# Selection for dispersal under larval malnutrition results in a non-monotonic kernel in *Drosophila melanogaster*

**DOI:** 10.1101/2023.10.26.564216

**Authors:** B. Vibishan, Akshay Malwade, Swaraj Ranjan Gupta, Anish Koner, Vinayak Khodake, Sutirth Dey

## Abstract

Dispersal is a key strategy for organisms to track favorable conditions and a shifting global climate has necessitated its closer investigation. While the distribution of dispersal distances (i.e. dispersal kernel) influences several aspects of population ecology, it remains unclear how kernel features are themselves affected by environmental factors. The current study addresses this question using laboratory populations of *Drosophila melanogaster* selected for greater dispersal under a protein-deficient larval diet. Our results reveal that dispersal-selected populations initiate dispersal more often and travel further on average than unselected controls. Although there is a significant change in the overall kernel shape, selection has not led to a greater tendency for long-distance dispersal (LDD). This is contrary to the results from a previous study on dispersal evolution under standard nutrition. We also find that dispersal evolution under larval malnutrition leads to a non-monotonic dispersal kernel, with the highest frequencies at intermediate distances. The evolved shape of the dispersal kernel and the limited LDD phenotype therefore point to significant qualitative shifts in the evolution of the dispersal kernel due to a change in larval nutrition. Our results thus provide a starting point for further investigation of such context dependence in dispersal evolution.

## 1 Introduction

Dispersal is one of the primary processes that enable organisms to track favorable conditions in space and time, making it an important consideration in the study of how organisms respond to rapid global climate change (Kokko and López-Sepulcre, 2006, Massot et al., 2008, Clobert et al., 2012). Although it is a complex, multi-component phenomenon, dispersal is usually divided into three distinct stages: (1) emigration/departure, (2) transfer or inter-patch movement, and (3) immigration and settlement (Bowler and Benton, 2005). Each stage involves specific environmental pressures, and the ability of the dispersing propagule to respond effectively to these varied pressures determines its eventual viability (Bonte et al., 2012).

In the departure phase, for instance, the tendency to leave the natal patch is often associated with selection to reduce kin competition (Hamilton and May, 1977), or avoid inbreeding (Bengtsson, 1978, Charlesworth and Charlesworth, 1987) and the effects of demographic stochasticity (McPeek and Holt, 1992, Travis and Dytham, 1998). Similarly, the settlement phase requires accurate assessments of patch quality and resource abundance (Bowler and Benton, 2005), as well as the ability to withstand competition from the natives of the new patch (Léna et al., 1998). Accordingly, organisms are known to respond to broader landscape features like the distribution of suitable habitats (Matter and Roland, 2002, Schneider et al., 2003, Entling et al., 2011), as well as the presence of conspecifics (Serrano and Tella, 2003, Clobert et al., 2009) in making decisions regarding leaving the natal patch or settling at a given patch. Furthermore, movement between patches is particularly vulnerable to a range of biotic and abiotic environmental influences that can be the causes of both mortality (DiLeo et al., 2022) as well as adaptive change (Travis and Dytham, 2002, Baines et al., 2020). Factors like temperature and resource availability can act as filters that pose extrinsic mortality risks and thus limit the spatial extent over which a population can viably spread (Denno et al., 1996). On the other hand, constraints during the movement phase could impose further requirements on dispersing individuals, including the development of physiological and morphological features, and behaviours (Zera and Denno, 1997, Bell et al., 2005). This would effectively determine which subset of a given population is able to carry out dispersal, and could affect the spatial segregation of dispersers and non-dispersers over a given landscape. Across its stages therefore, the process of dispersal is acted upon by a range of factors that establish the context in which dispersal occurs, all of which eventually determines the spatial distribution of the population.

An essential feature of dispersal that also offers a straightforward assessment of the spatial distribution of a population is the dispersal kernel (or the dispersal distance kernel), which is defined as the distribution of the frequency of propagules/individuals with distance from the source or a given natal patch (Clobert et al., 2012). The shape and range of the kernel are important features that dictate the speed of population spread under a range of patch distributions (Kot et al., 1996, Clark et al., 2001, Lindström et al., 2011), metapopulation structure and dynamics (Travis and Dytham, 1998, Higgins and Cain, 2002, Baines et al., 2020), population stability (Murrell et al., 2002), and phenotypic polymorphism (Travis and Dytham, 1999, Bonte et al., 2010). On the one hand, theoretical investigation using a variety of modelling frameworks has led to significant phenomenological and mechanistic understanding of the ecological and evolutionary consequences of the kernel features like shape and scale (Travis et al., 2010, Lindström et al., 2011). Moreover, while empirical measurements of dispersal are limited by logistic and practical considerations, field data over the years have nevertheless documented a variety of dispersal kernels across different study systems and ecological contexts (Clobert et al., 2012). On the other hand, studies investigating the potential for evolutionary and/or adaptive change in the dispersal kernel itself for a given population are relatively scarce. Heritable variation has been reliably identified in multiple aspects of the dispersal phenotype (Saastamoinen, 2008) and in select cases, the evolution of dispersal phenotypes has been demonstrated explicitly in the context of specific environmental changes (Stevens et al., 2012). Given that the dispersal kernel is a cumulative reflection of a large number of individual dispersal events, it is therefore reasonable to expect that adaptive change in the underlying dispersal traits could also lead to measurable shifts in the dispersal kernel. The dynamics of such shifts in the kernel and its potential context dependence remain under-explored.

Previous work has shown that selection for increased dispersal in large populations of the fruit fly, *Drosophila melanogaster* can lead to much more leptokurtic (fat-tailed) kernels than the corresponding control populations (Tung et al., 2018). More importantly, the selected populations in that study had a higher frequency of long-distance dispersers (LDDs), i.e., individuals that travel distances much further than the average of a population and are known to play a disproportionate role in the spatial spread and viability of a population (Petrovskii and Morozov, 2009). LDDs have profound impacts on several aspects of population dynamics and evolution. Both theoretical and empirical studies indicate that dispersal kernels with fatter tails, and thus higher frequencies of LDDs, lead to an accelerating invasion front in an expanding population (Kot et al., 1996, Andow, 2022). The shape of the kernel tail in particular has been shown to be relevant to dispersal-related responses to environmental disturbances (Bianchi et al., 2009). LDDs could therefore play a key role in how organisms respond to progressive habitat fragmentation and other sustained environmental shifts (Murrell et al., 2002). Higher absolute distances covered also relates to the spatial extent of a population (Murrell et al., 2002, Caton et al., 2022), which corresponds to the physical distances over which a population is seen to exist.

In this context, the distance over which an organism moves should depend on its physiological conditions and various behavioural patterns (Meylan et al., 2002, Clobert et al., 2009, Wijerathna and Evenden, 2019), which in turn will depend on various biotic and abiotic conditions in the natal patch as well as the movement landscape. For example, natal patch quality affects behavioural variability in the initiation of dispersal (Vinatier et al., 2011), as well as decisions related to exploration and settlement (Bonte et al., 2012, Jones et al., 2019). At the landscape level, spatial heterogeneity in resource availability could interact with the energetic costs of searching movements to make the evolution of optimal search strategies highly context-dependent (Zollner and Lima, 1999). At the individual level, the disperser’s nutritional status and the mobilization of existing energy reserves are both key determinants of the initiation and success of dispersal events (Saastamoinen and Rantala, 2013, Wijerathna and Evenden, 2019). All these factors can potentially lead to qualitatively different kinds of dispersal kernels under varying nutritional contexts. Does this mean that changing the food available to a population selected for dispersal can alter the shape of the evolved dispersal kernel?

In fruit flies specifically, it is well-known that larval nutritional status plays a crucial role in the amounts of energetic reserves available to the adult fly at eclosion (Vijendravarma et al., 2012), which would subsequently determine the fly’s capacity to actively travel long distances in the absence of food or water (Bonte et al., 2012). Larval nutrition is relatively easier to control over evolutionary timescales as well as single-generation experiments in laboratory populations of fruit flies (Prasad and Joshi, 2003, Tennessen et al., 2014, Tatar et al., 2014). Thus, *Drosophila melanogaster* offers a convenient system to investigate the context dependence of kernel evolution.

In this study, we select multiple populations of fruit flies for higher ambulatory dispersal under conditions of larval malnutrition (20% reduction of dietary protein, see S4 Text for details). The primary aim here is to look for signs of altered patterns of kernel evolution due to the additional pressure of larval malnutrition alongside dispersal selection. This involves measurements of the dispersal kernel in both selected and unselected populations across multiple generations of experimental evolution, allowing us to characterize several aspects of the dispersal phenotype, and follow their response to dispersal selection as well as larval malnutrition. The data arising out of these experiments reveal that larval nutrition places significant constraints on the extent and nature of dispersal evolution. In particular, while selection for dispersal under larval malnutrition leads to the evolution of a higher capacity for dispersal, the data shown here suggest that a concomitant tendency for long-distance dispersal is notably absent in the dispersal-selected populations. Moreover, control populations not selected for dispersal also showed an increase in some components of dispersal. The results of this study thus implicate natal nutrition as a major influence on kernel evolution and spatial spread of a population, and further substantiate the current understanding of the context dependence of dispersal evolution.

## 2 Materials and methods

### 2.1 Fly maintenance and selection protocol

The maintenance procedure of the selected and control flies were similar to previous studies (see Tung et al., 2018, and the supplementary material therein). Briefly, in the previous study, two populations had been derived from each of four ancestral outbred populations (DB_*i*_). The so-called VB_1−4_ populations were selected for higher ambulatory dispersal and the so-called VBC_1−4_ acted as the corresponding controls that did not undergo selection for dispersal. Populations that shared a subscript (for example, VB_1_ and VBC_1_) were related by ancestry (i.e. derived from DB_1_), and were treated as a block in statistical analyses.

In the current study, from each VBC_*i*_ (*i* ∈ [1 − 4]), we derived two more populations-MD_*i*_ (<u>M</u>alnourished <u>D</u>ispersers) and MC (<u>M</u>alnourished <u>C</u>ontrol). As detailed below, MDs were selected for higher ambulatory dispersal using a setup identical to that in the VB selection experiment, while MCs were maintained as corresponding unselected controls. Importantly, while VBs were selected for higher dispersal under standard nutrition, MDs underwent selection for higher dispersal under a larval malnourishment regime, which consisted of a modified banana-jaggery medium with one-third the amount of dry yeast powder as the standard recipe (S4 Text and Supplementary Table S1). This corresponds to a ∼ 20% reduction in the amount of protein, and initial observations showed that this leads to a slight decrease in egg-to-adult survivorship and an increase in the time to eclosion (personal observation). This is consistent with the known effects of increased larval crowding that have been demonstrated in the context of larval malnutrition, both by overall dilution of all dietary components as well as protein deficiency (Kolss et al., 2009, Lee et al., 2008).

Eggs from both MD and MC were reared until eclosion in this protein-deficient diet, at a density of about 350 eggs per 60mL of food in plastic milk bottles. The adult flies were subjected to dispersal selection on the 12th day after egg collection. The dispersal selection apparatus consisted of a set of two transparent plastic containers, one of which served as the source and the other as the sink. They were connected by transparent plastic piping (internal diameter, ∼ 1cm) that served as the dispersal path. The path extended to some distance inside the sink container to reduce back flow of flies from the sink. Neither the source nor the dispersal path contained food or water sources, while a strip of moist cotton was placed in the sink to prevent desiccation after dispersal. Starting from 2m in the first generation, the path length used for dispersal selection was increased intermittently across generations. This served as an approximation of progressively increasing habitat fragmentation, leading to a path length of 54m by the 93^*rd*^ generation when the last of the assays reported here were performed.

On day 12 from egg collection, adult MD flies of both sexes were introduced into the source container and allowed to migrate for a period of 6h, or until about 50% of the flies had reached the sink, whichever occurred first. In parallel, MC flies were introduced into plastic containers identical to the source containers of the dispersal setup that were not connected to a dispersal path. Halfway through dispersal selection, MC flies were given water through a moist piece of cotton, to mimic the desiccation conditions faced by the MD flies that only get water after successfully reaching the source. Following selection, MD flies that reached the sink and all the MC flies were transferred to Plexiglass population cages (25cm *×* 20cm *×* 15cm) with standard banana-jaggery medium. Yeast supplementation was provided on day 13 and a fresh food surface was provided overnight for oviposition on the evening of day 14. Eggs for the next generation were collected the following day, leading to a 15-day life cycle.

MD_*i*_-MC_*i*_ with the same subscript *i* were related by ancestry to the corresponding VBC_*i*_, were assayed together and considered a single block in statistical analyses. All populations were maintained at a breeding population size of roughly 2400 flies, in discrete generation cycles under uniform environmental conditions comprising a temperature of 25^°^C and 24h light conditions. All assays were performed after one generation of common garden rearing under the larval malnourishment regime, but without dispersal selection, to minimize the effects of non-genetic parental effects due to dispersal selection. In order to avoid larval crowding, eggs for pre-assay common garden rearing were collected at a density of about 350 per bottle containing about 60mL of food, and for the assays, about 30 eggs per vial containing about 6mL of food.

### 2.2 Kernel experiments and calculations

For the purposes of this study, the dispersal kernel was measured as the number of flies travelling up to a certain distance along a one-dimensional path within a given time period. This corresponds to the accepted definition of a dispersal distance kernel across a variety of study systems (Clobert et al., 2012). Kernels were measured thrice over the period of selection, at generations 42, 66-67 (blocks 1 and 2 in 66, and 3 and 4 in 67) and 93. For each kernel, a source-path-destination setup similar to that used for dispersal selection was assembled, consisting of detachable segments. A 20m path was used in generation 42 with a combination of 0.5m and 1m segments, and a 60m path was used in generation 66-67 comprising only of 1m segments. In keeping with the selection protocol, on day 12 from egg collection, a mixed-sex cohort of about 1000 adult flies of the respective population were introduced into the source and allowed to disperse for 6h. In generations 42 and 66-67, no food was provided in the source container and no water was provided in the destination container. At the end of 6h, the dispersal setup was disassembled and each individual segment was sealed before being placed in a hot air oven at ∼ 40^°^C for about 30 minutes to heat-kill the flies, after which the numbers of male and female flies in each segment were recorded. All the flies from a given segment were considered to have travelled the same distance, calculated as the distance from the source up to the midpoint of the corresponding segment. The 93^*rd*^ generation kernel was performed with a 2m path and no water in the destination container, but included about 25mL of standard banana jaggery food in the source container. The goal of this shorter kernel was to test whether or not dispersal evolution under larval malnutrition resulted in phenotype-dependent dispersal. Since the proportion of flies leaving the source container in presence of food is a sufficient measure for this purpose, a shorter path length was considered appropriate. Three replicate kernels were run for each population in generations 42 and 93, and two replicates were run in generation 66-67, and all of them were conducted two blocks at a time due to logistic constraints.

The above data were used to derive the following characteristic quantities that together provide a simple description of the dispersal kernel: (1) dispersal propensity, calculated as the proportion of total flies that left the source container by the end of the 6h period; (2) dispersal ability, which represents the average distance travelled by the flies of a given population and was calculated as 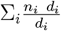, where *n*_*i*_ is the number of flies in the *i*th segment and *d*_*i*_ is the distance from the source to the midpoint of the *i*th segment. Note that ability was only calculated for flies that left the source container in order to prevent biases arising from different frequencies of flies staying in the source between the two populations; (3) Higher moments including the standard deviation, skew and kurtosis, to describe the overall shape and symmetry of the kernel; and (4) The 90^*th*^ percentile of the distribution of distances travelled, which gives an indication of the spatial extent of the kernel. The shorter kernel from generation 93 produced only sex-separated estimates of dispersal propensity for each population. Calculations were performed for each block separately, averaging over within-block replicates.

Nonlinear regression was used to further characterize any divergence in the kernel properties between selected and control populations by fitting a curve to the distribution of distances travelled. While the negative-exponential function has been a popular choice for insect dispersal kernels (Taylor, 1978, Hill et al., 1996, Clobert et al., 2012), visual inspection of our data suggested that the MD kernel was unimodal while the MC kernel seemed to decay in a more prototypical monotonic decline. The Gamma distribution can mathematically produce both unimodal peaks and monotonic declines, and has been applied to a variety of dispersal-related processes (Van Houtan et al., 2007, Guttal et al., 2011). The following form was used to fit the distribution of frequency of flies with distance:

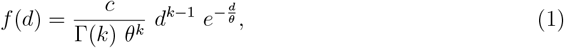

where *f* (*d*) is the frequency of flies (ranged [0, 1]) at a distance *d* from the source. This form included three free parameters whose values must be determined by optimization: *θ*, the horizontal scale parameter, *k*, the shape parameter, and *c*, an additional multiplicative factor introduced by us to scale the distribution along the vertical axis (Van Houtan et al., 2007, Clobert et al., 2012). As with dispersal ability, the source compartment was ignored for curve fitting. Nonlinear least-squares regression was carried out to fit Equation 1 to kernel data from generations 42 and 66-67 using the `nls multstart()` function from the `nls.multstart` package (Padfield and Matheson, 2020, version 1.2.0) in R (R Core Team, 2021, version 4.1.2). Since convergence of a nonlinear regression is highly sensitive to the starting values of the free parameters, `nls` `multstart()` was used to run the regression with 500 initial values picked randomly from a given range and the best model was chosen based on AIC (Akaike information criteria) values. Visual inspection of plots of the residuals did not reveal any significant biases in any of the fits, except for the tail of the MC kernel where zero inflation might have contributed to some overestimation towards the end of the kernel (not shown). The Gamma function was fit to frequencies averaged across blocks for the illustrative plots in Figure 4, while S1.6 Text and S2.6 Text give the fitted parameter values of the Gamma function as well as other statistical information, separated by block and sex.

As mentioned earlier, long-distance dispersal is primarily studied in the context of individuals that move significantly beyond the average dispersal distance out of a given population. Keeping with this conceptual definition, differences in long-distance dispersal between MD and MC were characterized based on the extreme parts of the dispersal kernel using two approaches. Hampel filters are commonly used to identify data points that could be considered outliers based on the median absolute deviation and a fixed cutoff value, and are particularly preferred for being agnostic to the exact shape of the underlying distribution (Wilcox, 2010). The frequency of extreme dispersal events could therefore be reflected in the number of outliers identified by the Hampel filter. For the first approach to characterize LDD therefore, outlier identification was carried out with a custom `R` script for a Hampel filter with a cutoff value of 2.5 (Wilcox, 2010) and the number of outliers detected was examined in the MD and MC kernels. In the second approach to LDD, the aim was to identify whether or not an observed change in the extreme part of the kernel is equivalent to an underlying change in the average of the kernel, or is incommensurate with the latter. To do this, the average of distances above the 95^*th*^ distance percentile was calculated for both MD and MC-this represents the absolute magnitude of the top 5% of all dispersers. This extreme value was then scaled by the corresponding overall average distance, to normalize the extreme value against the mean. If selection for dispersal in MD has led to a higher proportion of long-distance dispersers (LDDs) in addition to greater dispersal in general, this scaled extreme value of the kernel would be higher for MD than MC, in excess of any underlying difference in the average distance moved.

### 2.3 Statistical analyses

Unless specified otherwise, all analyses were performed over block-segregated, sex-separated mean values of the response variable, averaging only over the measurement replicates within each block. Data from different generations were analyzed separately such that generation was never included as a factor in any part of the analysis. As detailed below, all models were fit to averages over measurement replicates within a block, and data were never averaged across blocks or sex in any of the analysis, except for the demonstrative plot of the Gamma-fitted dispersal kernels in Figure 4. All analyses were carried out on the `R` platform (R Core Team, 2021, version 4.1.2).

For dispersal propensity, mean numbers of flies that left the source and those that stayed in the source were used as “Success” and “Fail” values in a mixed-effects GLM with binomial error distributions and logit link function, with selection and sex as fixed effects, and block as a random intercept. Mean values of dispersal ability were fit to a Gaussian mixed-effects GLM with selection and sex as fixed effects, and block as a random intercept. Both GLMs were fit using the `lmerTest` package (Kuznetsova et al., 2017, version 3.1-3), and functions from the `emmeans` (Lenth, 2023, version 1.8.4-1) and `psych` (Revelle, 2022, version 2.2.9) packages were used to calculate marginal differences between group means and effect sizes respectively for pairwise comparisons. For the latter, values of Cohen’s *d* were interpreted as large, medium and small for |*d*| = 0.8, 0.8 *>* |*d*| *>* 0.5, and |*d*| *<* 0.5 respectively (Cohen, 2013). Higher moments of the kernel including SD, skew and kurtosis were compared across the four blocks between MD and MC using Wilcoxon’s rank-sum tests using the `wilcox.test()` function from the `stats` package (R Core Team, 2021, version 4.1.2).

The procedure for nonlinear regression of the Gamma distribution function has been elaborated earlier. Goodness of fit of the Gamma distribution was also tested using *χ*^2^ and Kolmogorov-Smirnov tests using the `chisq.test()` and `ks.test()`functions from the `stats` package. Fitted values of the Gamma distribution parameters, *θ, k* and *c* were used to compare the properties of the kernel. For an unscaled Gamma distribution, the mean and SD can be expressed as *kθ* and 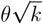 respectively. From Equation 1, the vertical scaling factor *c* can be used to derive a corrected value of *θ* that corresponds to the canonical scale parameter such that, 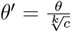. The mean and SD for the scaled Gamma distribution in Equation 1 can therefore be calculated as *kθ*^*′*^ and 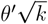 respectively. *k* and *θ*^*′*^ can be meaningfully compared between MD and MC, for each of which, Gaussian mixed-effects GLMs were used with selection and sex as fixed effects, and block as a random intercept. Sexual dimorphism in overall kernel shape was investigated using population-wise block-segregated Kolmogorov-Smirnov tests to test the null hypothesis that the male and female kernels are derived from the same distribution.

The unscaled and scaled means of distances above the 95^*th*^ percentile were compared across the four blocks between MD and MC using Wilcoxon’s rank-sum tests using the `wilcox.test()` function from the `stats` package.

In all cases, the main results are reported with appropriate statistical information in brief. The full results of all statistical procedures have been given in S1, S2, and S3 Text for generations 42, 66-67, and 93 respectively.

## 3 Results

### 3.1 Dispersal components respond to selection under larval malnutrition

The flies selected for dispersal (i.e. MDs) had greater propensity compared to the corresponding controls (i.e. MCs) (Figure 1), with the selection *×* sex interaction being significant at both generation 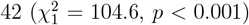 and 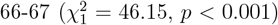. Pairwise comparisons in generation 42 showed that while MD propensity was higher than that of MC, the difference among females was much greater than that among males (MD vs. MC, Cohen’s *d* = 9.4 for females vs. 2.8 for males; S1.1 Text). Data from the 93^*rd*^ generation further show that MC propensity is condition-dependent, and drops to its lowest value in the presence of food even as MD propensity remains largely unchanged (Figure 1C, see S3 Text for details). This indicates that while dispersal propensity in MC flies is affected by the presence of food, propensity in MD flies is constitutively high regardless of the presence or absence of food.

**Figure 1.**
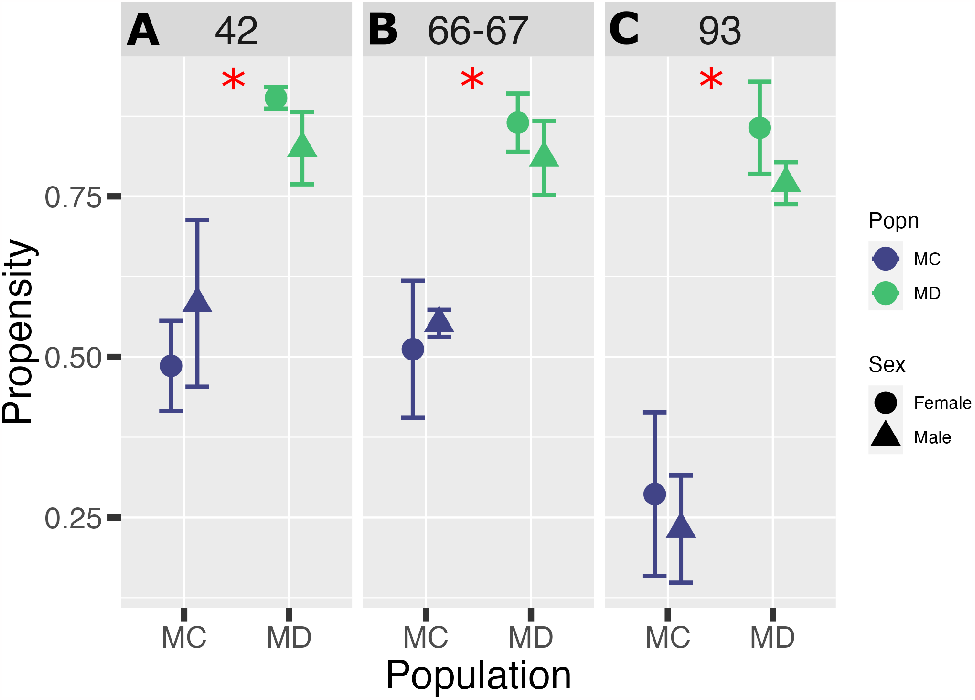
Increase in dispersal propensity due to dispersal selection. Assays for generations (A) 42 and (B) 66-67 were performed without food in the source while generation (C) 93 had food in the source. All points are ensemble means *±* SD over the four blocks. * denotes large effect difference (see S1.1 and S2.1 Text for details).

Figure 2 shows that on an average, the MDs traveled a longer distance compared to the MCs (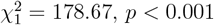 in generation 66-67). Comparing trends across generations, it is clear that there is a nearly three-fold increase in ability between the two generations not only in MDs, but also MCs, despite the latter not being selected for dispersal (*cf* Figures 2A and B; see Discussion for a possible reason for the latter observation). A notable consequence of this increase is that the dispersal ability of MC at generations 66-67 has in fact exceeded that of MD from generation 42.

**Figure 2.**
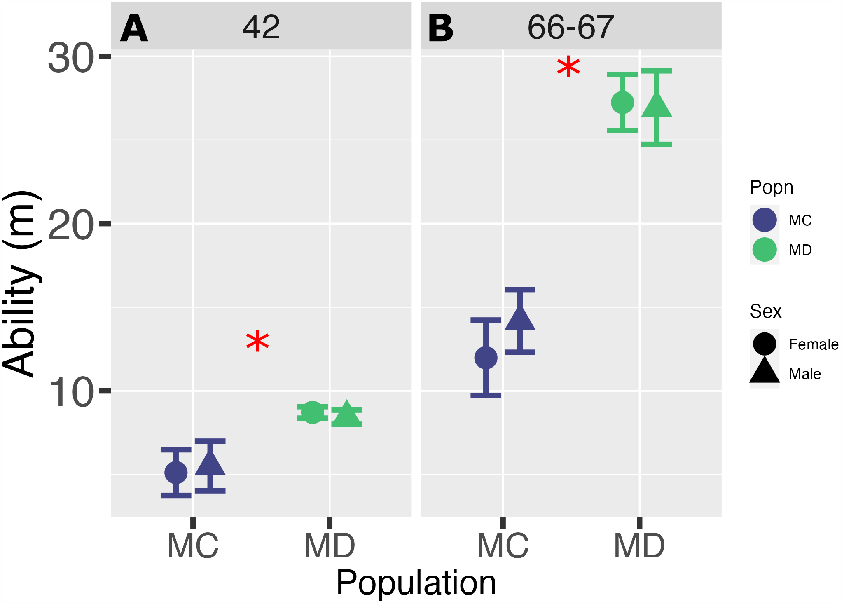
Progressive increase in dispersal ability. Data only from generations (A) 42 and (B) 66-67 as dispersal ability was not measured in generation 93. All points are ensemble means*±*SD over the four blocks. * denotes large effect difference (see S1.2 and S2.2 Text for details).

### 3.2 Kernel shape progressively diverges over ∼ 20 generations of selection

As Figure 2 shows, between generations 42 to 66-67, average dispersal ability had increased considerably, in both MC and MD. This suggests that the shape of the overall dispersal kernel could have likewise shifted, both between MD and MC, and over the course of 20 generations of selection.

Figure 3 presents a comparison of multiple shape characteristics of the dispersal kernel in terms of the higher moments of the distribution between MD and MC, in generations 42 (Figure 3A) and 66-67 (Figure 3B). Figure 4 shows the actual change in overall kernel shape over the same period. Between MD and MC, in both generations, the MD kernel is associated with lower skew and kurtosis than MC (Figures 3B and C, both columns), while a trend (not statistically significant) is seen towards higher standard deviation in MD than MC (Figure 3A, S1.3 and S2.3 Text). In both generations, MC specifically has a considerable right skew whereas MD kernels are more symmetric. Kurtosis is calculated here as the absolute value, so that a kurtosis value of 3 corresponds to a normal distribution. The data suggest that the MD kernel, although more apparently bell-shaped, is rather platykurtic as its tails decay faster than those of a normal distribution. Although the moment values between the two generations cannot be compared within a single statistical framework, visual examination of the two columns in Figure 3 indicates that both the standard deviation and the 90^*th*^ distance percentile of both MD and MC kernels have increased. This is consistent with the trends in average dispersal ability in Figure 2, and suggests that over the course of 20 generations of selection, both populations have developed the capacity to spread over much greater physical distances, i.e. increased their spatial extent (*sensu* Murrell et al., 2002).

**Figure 3.**
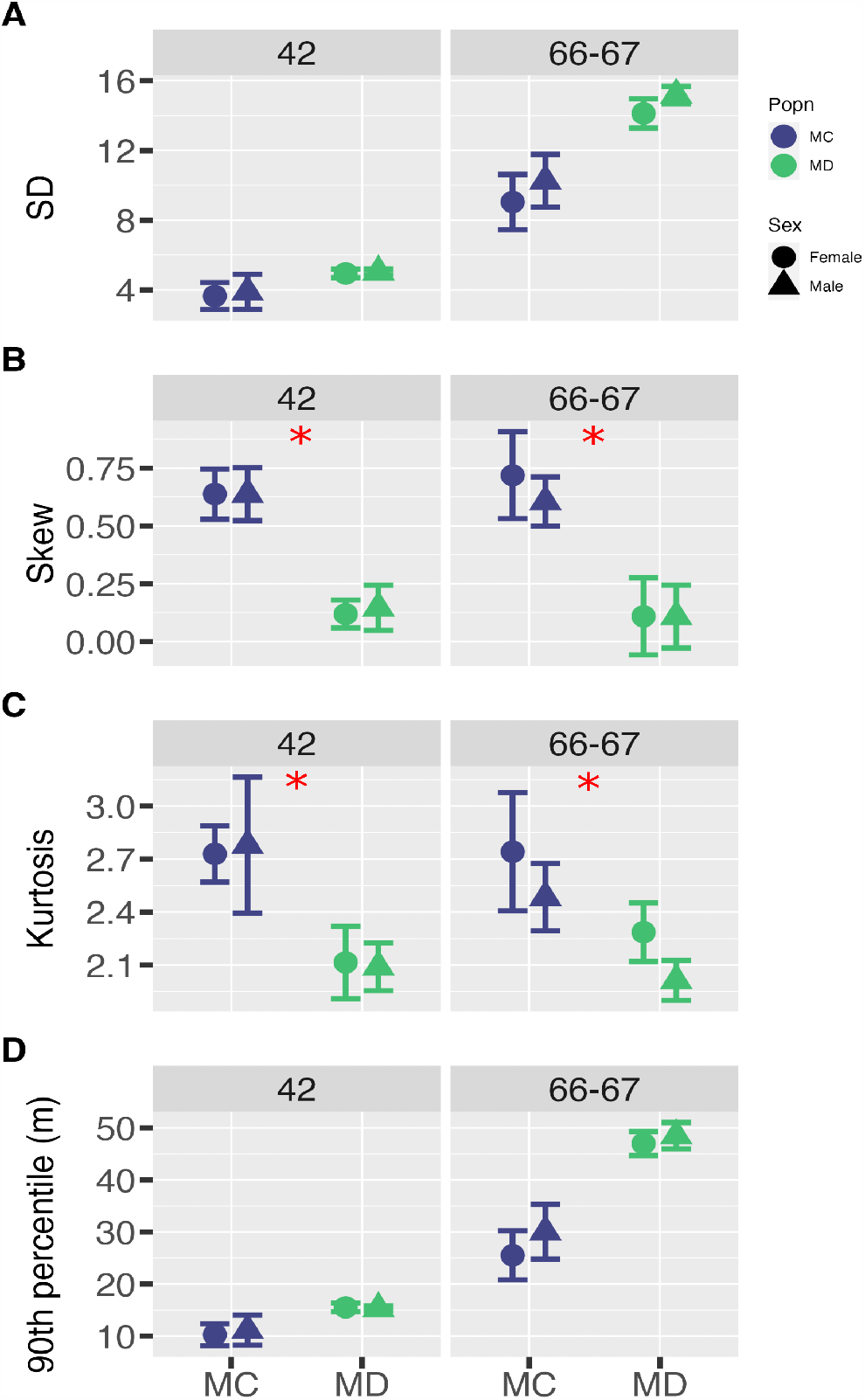
Model-free estimates of the dispersal kernel shape. Higher moments of the dispersal kernel, including (A) the standard deviation (SD), (B) skew, and (C) absolute kurtosis, along with (D) the 90^*th*^ percentile of the absolute distance moved by each population. All points are ensemble means*±*SD over the four blocks. * *p <* 0.05, Wilcoxon’s rank-sum test (see S1.3 and S2.3 Text for details).

**Figure 4.**
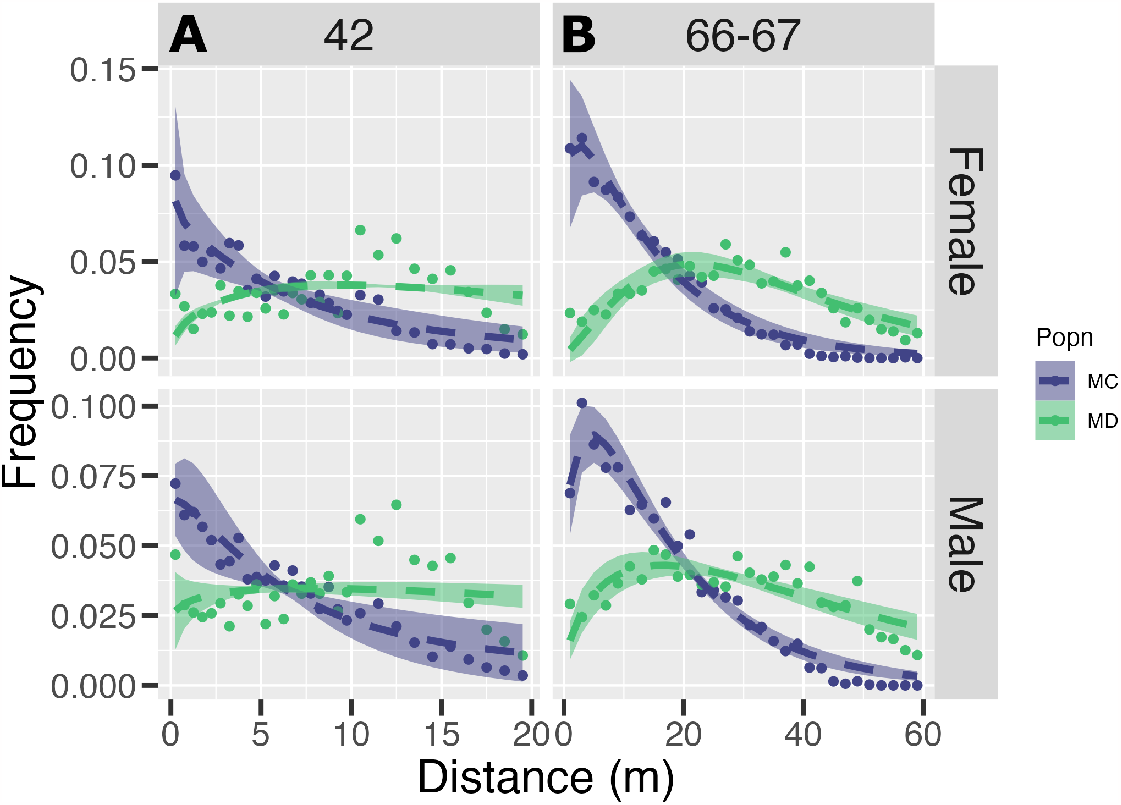
Gamma function as given in Equation 1 fit to the distribution of frequencies with distance. The points are mean observed frequencies across the four blocks, the dashed lines are corresponding predicted values based on the fit and the shaded region is the standard deviation of the predicted values, all calculated as ensembles over the four blocks. Goodness of fit was assessed using Kolmogorov-Smirnov and *χ*^2^ tests to compare the observed frequency with model predictions (see S1.4 and S2.4 Text for goodness of fit, and S1.6 and S2.6 Text for a list of the fit parameter and AIC values, separated by block and sex).

The differences seen in the higher moments between both MD and MC, as well as generations 42 and 66-67, are reflected in the shapes of the MC and MD kernels. The MD frequency distribution stretches much further to the right than the MC distribution, and that from generation 42 to 66-67, the physical length of both kernels have increased (Figure 4, note the different x-axis limits between 4A and B). The fitted values of the Gamma distribution parameters are consistent with this change in shape, as both *k* and *θ*^*′*^ are on average smaller for MC than MD (Supplementary Figure S3, S1.6 and S2.6 Text). A comparison between generations 42 and 66-67 (*cf* Figures 4A and B) shows that the overall divergence between MD and MC has progressively increased over evolutionary time such that the difference in kernel shape between MD and MC is much clearer at generations 66-67 than at generation 42. The fact that the MD population kernels are clearly non-monotonic is noteworthy as empirical insect kernels in literature have been predominantly monotonic declines described by negative exponential or Gaussian tails (Koch et al., 2012).

### 3.3 Incipient sexual dimorphism in a subset of dispersal components

Among the various quantities derived from the dispersal kernel, sexual dimorphism in the difference between MD and MC is apparent in propensity while a trend may be emerging in ability (Figures 1 and 2). The selection *×* sex interaction for propensity is significant in both generation 42 and 66-67 (S1.1 and S2.1 Text). It is clear from the corresponding odds ratios that in MD, male propensity is lower than that of females, while the trend is reversed in MC. A similar trend is also indicated for ability, although neither the effects from the GLMM nor the differences between pairwise means is statistically significant (S1.2 and S2.2 Text). None of the higher moments calculated from the frequency distribution showed evidence of sexual dimorphism in either generation (S1.3 and S2.3 Text).

With respect to the Gamma functions fitted to the dispersal kernels, Supplementary Figure S3 suggests that differences in *θ*^*′*^ and *k* between MD and MC could be sex-specific. On the other hand, block-wise Kolmogorov-Smirnov tests revealed no significant differences in the overall shape of the kernel between males and females in either population (S1.5 and S2.5 Text).

### 3.4 Dispersal selection under larval malnutrition does not lead to higher LDD

Earlier work showed that populations selected for dispersal under standard nutrition had a considerable number of individuals that dispersed much further than the population average (long-distance dispersers or LDDs), at higher frequencies than the corresponding controls (Tung et al., 2018). Addressing this possibility in the current populations, a Hampel filter with a cutoff value of 2.5 failed to identify any outliers at all in the MD kernel while a varying but nonzero number of outliers were identified in MC (Figure 5A). This suggests that the overall frequency of long-distance dispersal events could be lower in MD than in MC. The unscaled mean of the top 5% distances travelled is significantly higher for MD than MC (Figure 5B, S2.7 Text), which is consistent with the existing difference in the average ability (Figure 2B). However, when scaled by the overall mean, the top 5% of distances travelled by MC flies are significantly higher than those of MD (Figure 5C, S2.7 Text). This is a further indication of the “long-tail” of the MC kernel, as MC flies travel greater distances relative to their mean than MD flies do relative to theirs. The data in Figure 5 altogether indicate that the MD kernel is composed of fewer long-distance dispersal events, and while the distances travelled by MD flies are on average much higher than MC, the latter are more extreme dispersers. Finally, Supplementary Figure S1 shows an alternate way of making the same comparison, based on the frequency distribution of all distances after having the corresponding mean subtracted from the entire distribution. Comparing the percentiles of mean-subtracted distances between MD and MC, no clear difference emerges at long distances from the source. In fact, towards the right hand extreme, the percentiles of MD and MC converge, indicating that the shapes of the kernel tails are very similar after accounting for the difference in distance moved. Taken together, these data suggest that while average dispersal has evolved as a result of selection in MD, long-distance dispersal has not.

**Figure 5.**
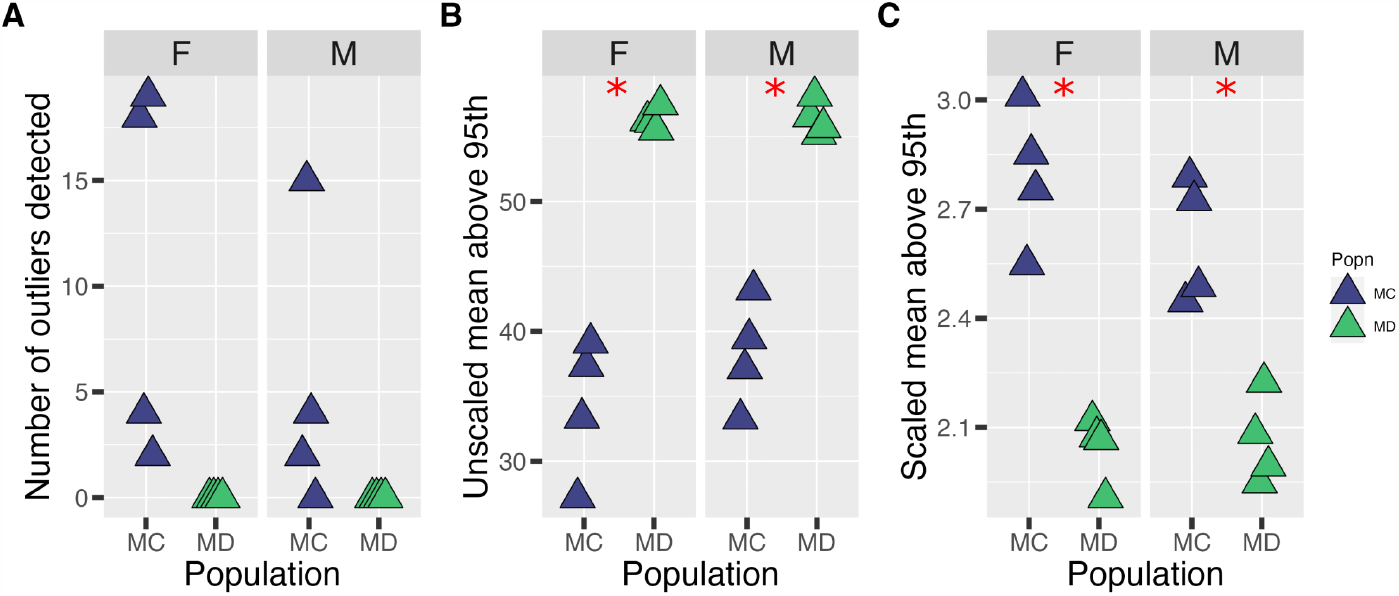
Measures of long-distance dispersal. (A) Number of outliers inferred using the Hampel filter with a cutoff value of 2.5; (B) Mean of absolute distances travelled above the 95^*th*^ percentile; and (C) mean of distances travelled above the 95^*th*^ percentile scaled by the mean distance over the entire kernel only for generation 66-67; * *p <* 0.05 by Wilcoxon’s rank sum test. Trends in generation 42 are qualitatively the same and shown in Supplementary Figure S2, with detailed statistical information in S1.7 and S2.7 Text.

## 4 Discussion

This study intended to examine how the dispersal kernel would evolve if organisms were subjected to a diet with relatively less protein content (compared to a standard laboratory diet). Several insights emerge from the data presented so far.

### 4.2 Larval malnutrition decouples short- and long-distance dispersal under selection

We find that selection for dispersal can lead to increases in both dispersal propensity and ability, as shown consistently by the MD flies (Figures 1 and 2). Furthermore, the increase in dispersal propensity is also seen to be phenotype-dependent, as it is not affected by changing the availability of food at the source. Both of these outcomes are consistent with the results of dispersal selection under standard nutrition (Tung et al., 2018), and it is therefore difficult to pinpoint a clear effect of the nutritional environment of dispersal selection based on these results alone. However, a closer look at the extreme parts of both the MD and MC kernels shows that while MD flies move greater distances on average, the frequency of LDDs-individuals moving much further than the average-is higher in MC than in MD (Figure 5, S1.7 and S2.7 Text). This is a key departure from previously-reported data on dispersal selection under standard nutrition (Tung et al., 2018), which showed that short-distance dispersal, as reflected by the average ability, increased concomitantly with long-distance dispersal. The pattern of kernel evolution seen here therefore suggests that the addition of larval malnutrition to the selection regime has effectively separated short- and long-distance dispersal.

Individual-based modelling (IBM) has previously suggested that landscape features such as habitat availability and fragmentation can lead to independent evolution of short- and long-distance dispersal (Bonte et al., 2010). Specifically, it was found that both increased habitat availability and evenly-distributed habitats favor the evolution of long-distance dispersal, whereas increasing habitat fragmentation and lower habitat availability generally discourages long-distance dispersal. In this context, it is relevant to note that both the current and the previous study on dispersal selection (Tung et al., 2018) involve progressively increasing path lengths in a reflection of habitat fragmentation. Taken together, these two studies indicate that while long-distance dispersal can evolve under conditions of habitat fragmentation under standard nutrition (Tung et al., 2018), the extent of long-distance dispersal is clearly constrained by severity of nutrient limitation in the larval and adult stages preceding dispersal. It is thus clear that theoretical expectations of dispersal evolution under changing landscape regimes, albeit a valuable starting point, need closer consideration of local ecological factors that can attenuate these effects. An indication of an attenuation of this kind can be seen in the documented effects of poor larval nutrition on LDD in our experimental system. We speculate that dispersal evolution under nutrient limitation has brought MD flies closer to their intrinsic physiological capacity for dispersal, thus impeding the evolution of long-distance dispersal (Figures 5B and C).

The current study thus documents a nutrition-driven separation of components of the kernel, which in turn is consistent with the broader dispersal literature. For example, meta-analyses of dispersal data from mammalian taxa (Whitmee and Orme, 2013) have identified that the predictive power of a wide range of ecological, life history and phylogenetic variables differs considerably between median and maximum dispersal distances, indicating that different parts of the dispersal kernel may be responding to different environmental and physiological factors. Disparate changes in dispersal components has been also documented empirically in butterflies, in which oviposition strategies and other routine movements correlated more strongly with short-distance dispersal (Hovestadt et al., 2011, Stevens et al., 2012). The insights from our study could therefore inform dispersal ecology beyond the laboratory settings of the fly system.

### 4.2 Larval malnutrition drives higher dispersal in the absence of dispersal selection

Across taxa and ecological contexts, natal patch conditions has been shown to affect the overall tendency of organisms to initiate and carry out dispersal, as the tendency to leave is known to be associated both with declining quality of the natal patch as well as higher frequency of competitors within the natal patch (Muller-Landau et al., 2003, Fragoso et al., 2003, Bowler and Benton, 2005, Latty and Reid, 2010, DiLeo et al., 2022). Trends in the dispersal characteristics of MC between generations 42 and 66-67 offer further indications of the ecological role of natal nutrition. Our data show that under larval malnutrition, MC flies have developed a marked increase in both dispersal propensity (Figure 1) and ability (Figure 2) despite not being selected for dispersal. It is also worth pointing out that propensity and ability could be independent components of dispersal and as such, affect different aspects of the dispersal process (Fronhofer et al., 2014, Bonte et al., 2012). Interestingly, a recent study has demonstrated that fruit flies under larval malnutrition through protein reduction show increased activity of certain dopamine signaling pathways leading to higher locomotor activity in a dopamine receptor-dependent manner (Zúñiga-Hernández et al., 2023). Therefore, it is possible that the larval malnutrition experienced by the MC flies had led to selection for increased levels of dopamine, which in turn could have increased their locomotor activity across generations. Although it is not possible for us to speculate further on this aspect based on the current data, it is a promising line for future investigation.

### 4.3 Insect dispersal kernels can be non-monotonic

Greater average dispersal (Figure 2) and limited long-distance dispersal (Figure 5) among the MDs together could cause the corresponding kernel to evolve towards a bell-shaped distribution, such that the frequencies of organisms are highest at intermediate distances from the source. On the other hand, the MC kernel retains its overall declining trend despite increasing in spatial extent (Figure 4). Prior theoretical work has shown that when the costs of dispersal increase linearly with distance travelled, dispersal distributions could evolve to show peak probabilities of dispersal at intermediate distances from the source (Rousset and Gandon, 2002), i.e., non-monotonocity. Our selection apparatus requires dispersal to occur under desiccation (Mishra et al., 2022), which could form the basis of increasing costs with dispersal distance, although it is difficult to say whether or not this increase is likely to be linear. Such increasing costs could be exacerbated by larval malnutrition, suggesting further possibilities for the evolution of non-monotonic dispersal kernels.

Non-monotonic kernels entail an uneven distribution of individuals across space, which could in turn lead to further spatio-temporal variability in the biotic and abiotic conditions experienced by the dispersers. While the evolution of dispersal is known to be impacted by environmental variability (Denno et al., 1996), spatial structure (Travis et al., 2010) and presence of other organisms (Muller-Landau et al., 2003) individually, the possibility of multiple such factors being tied together by the shape of the dispersal kernel is relatively unexplored, particularly at short to intermediate distances from the natal patch. It could thus be of interest to examine whether and how non-monotonic dispersal kernels can enforce spatial structures in ecological interactions, and what its consequences are for population ecology in general.

## S1 Text for

Statistical analyses for Generation 42

### S1.1 Propensity

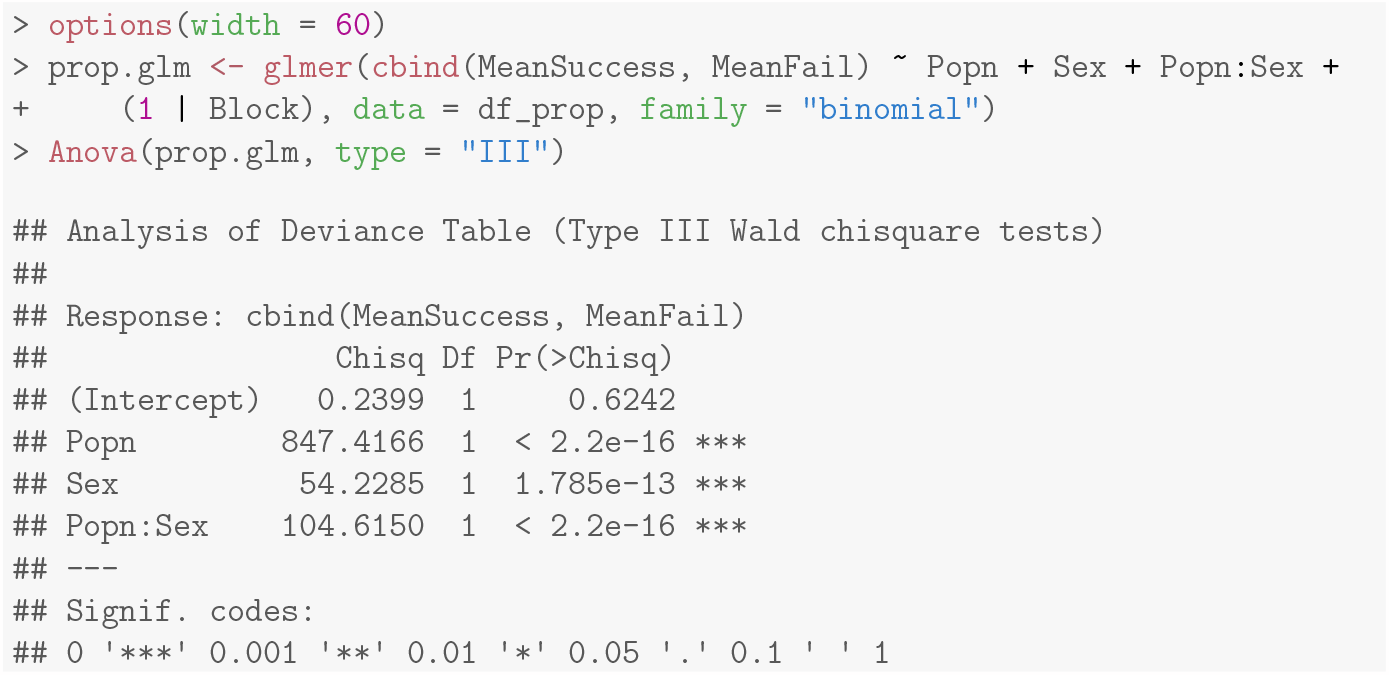

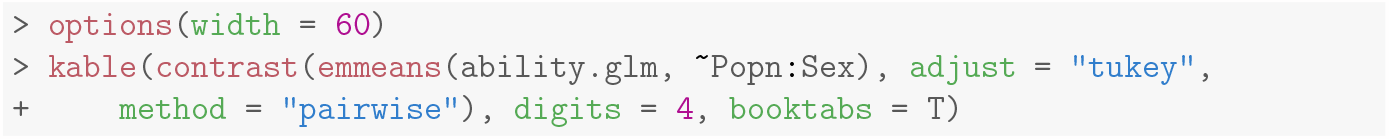

| contrast | odds.ratio | SE | df | null | z.ratio | p.value |
| --- | --- | --- | --- | --- | --- | --- |
| MC F / MD F | 0.0937 | 0.0076 | Inf | 1 | -29.1104 | 0 |
| MC F / MC M | 0.6686 | 0.0366 | Inf | 1 | -7.3640 | 0 |
| MC F / MD M | 0.1844 | 0.0124 | Inf | 1 | -25.0991 | 0 |
| MD F / MC M | 7.1328 | 0.5816 | Inf | 1 | 24.0934 | 0 |
| MD F / MD M | 1.9673 | 0.1776 | Inf | 1 | 7.4979 | 0 |
| MC M / MD M | 0.2758 | 0.0187 | Inf | 1 | -19.0447 | 0 |

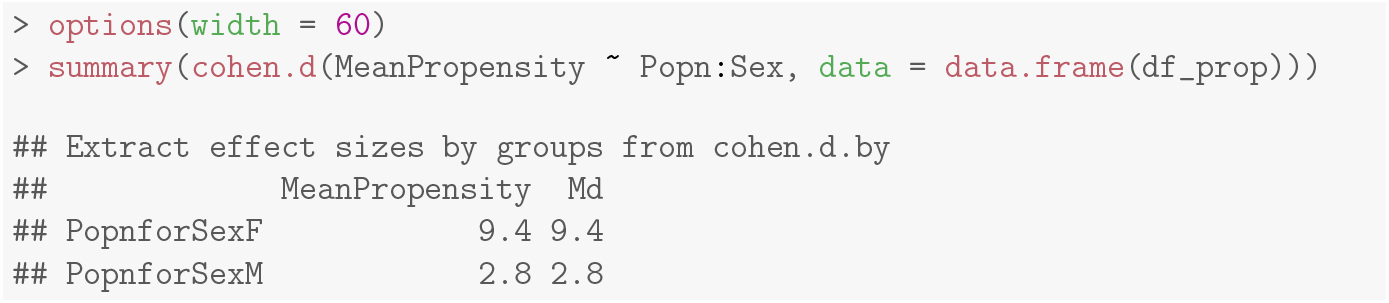

### S1.2 Ability

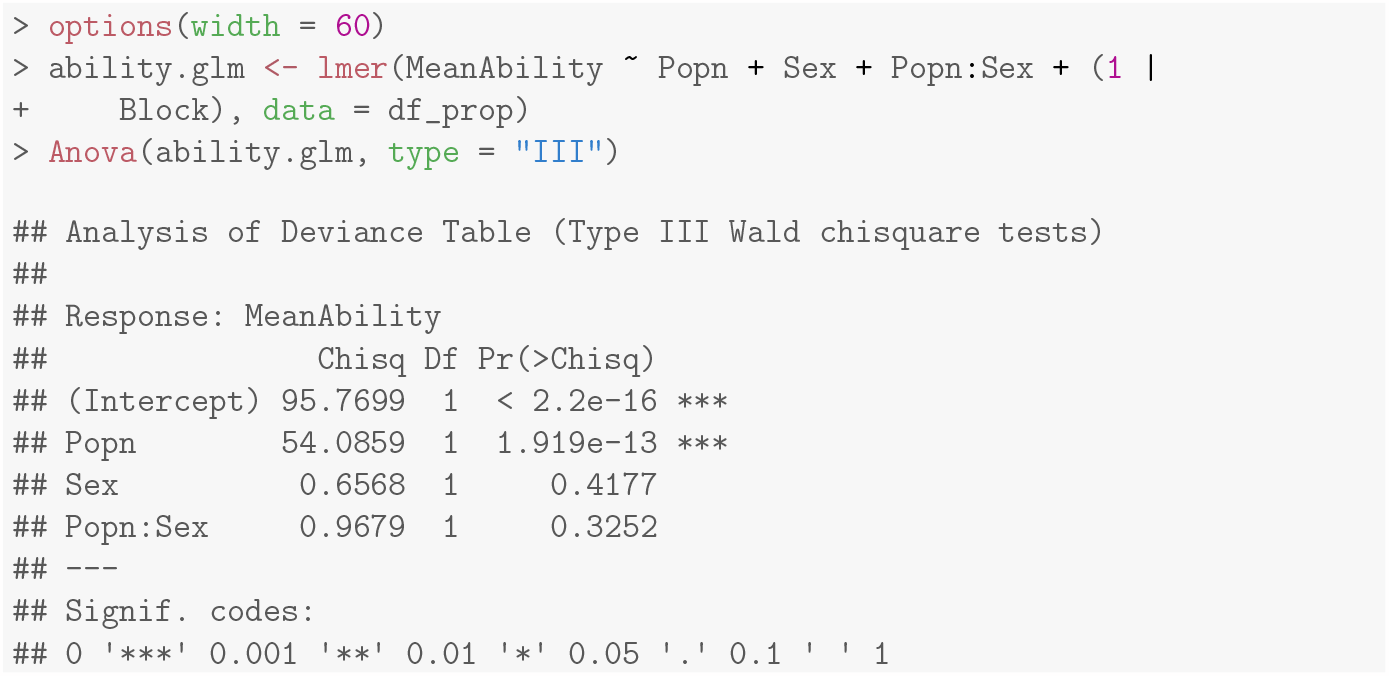

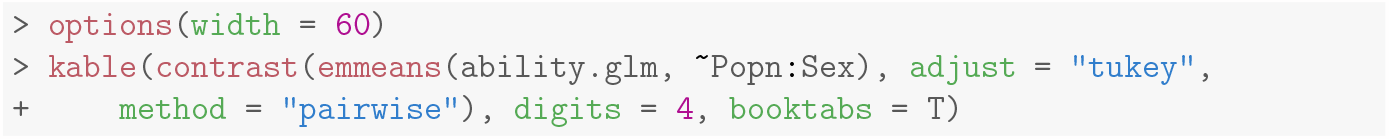

| contrast | estimate | SE | df | t.ratio | p.value |
| --- | --- | --- | --- | --- | --- |
| MC F - MD F | -3.6001 | 0.4895 | 9 | -7.3543 | 0.0002 |
| MC F - MC M | -0.3967 | 0.4895 | 9 | -0.8104 | 0.8481 |
| MC F - MD M | -3.3158 | 0.4895 | 9 | -6.7734 | 0.0004 |
| MD F - MC M | 3.2034 | 0.4895 | 9 | 6.5439 | 0.0005 |
| MD F - MD M | 0.2843 | 0.4895 | 9 | 0.5809 | 0.9354 |
| MC M - MD M | -2.9190 | 0.4895 | 9 | -5.9630 | 0.0010 |

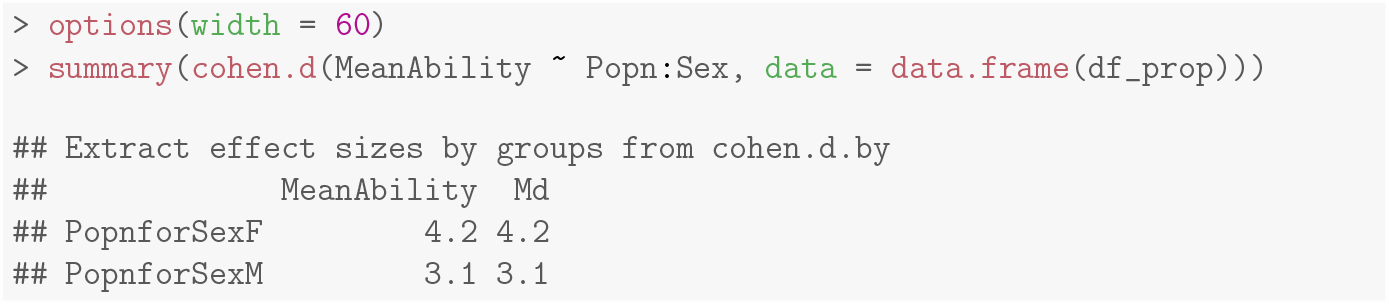

### S1.3 Higher moments of the kernel and 90th percentile

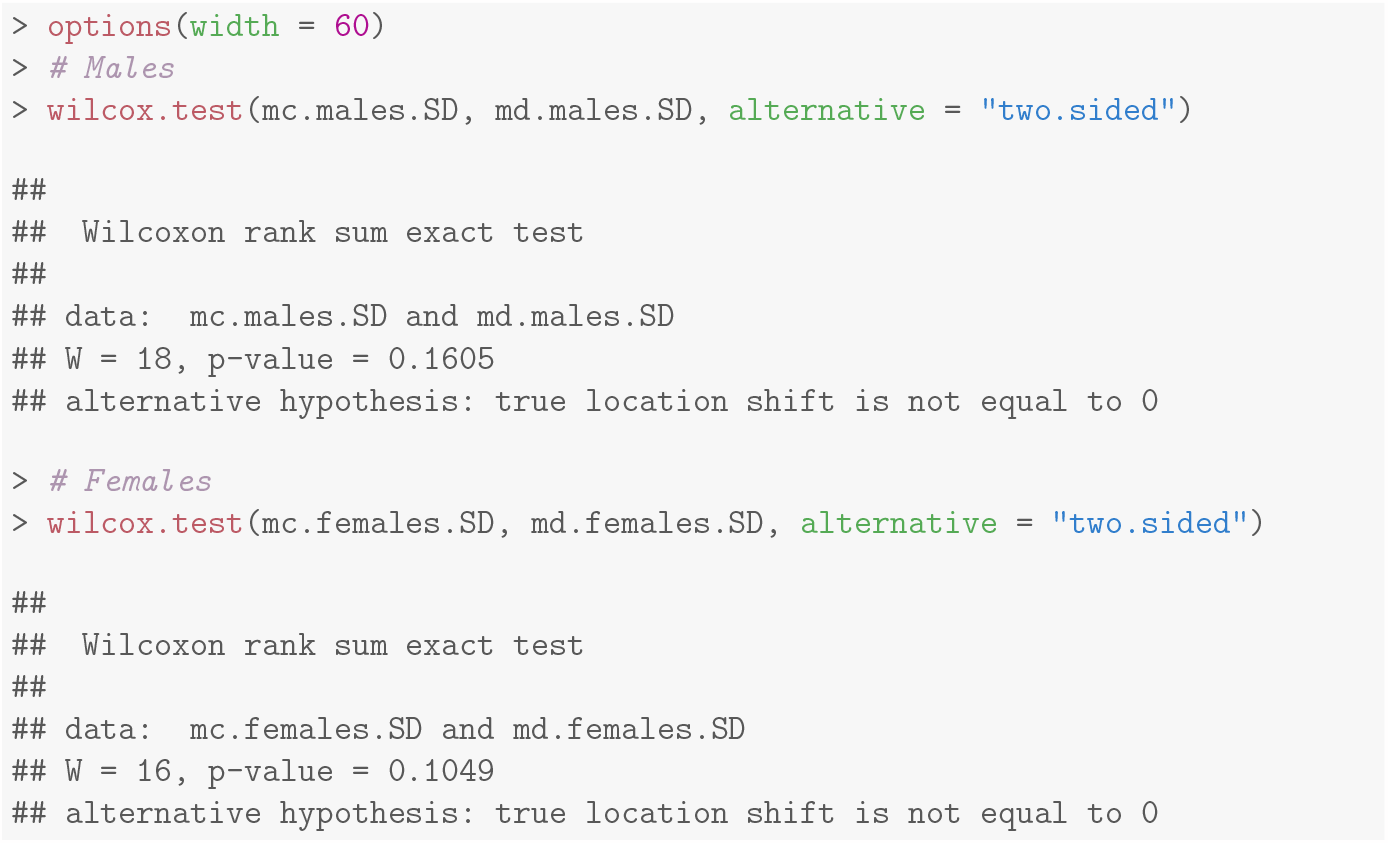

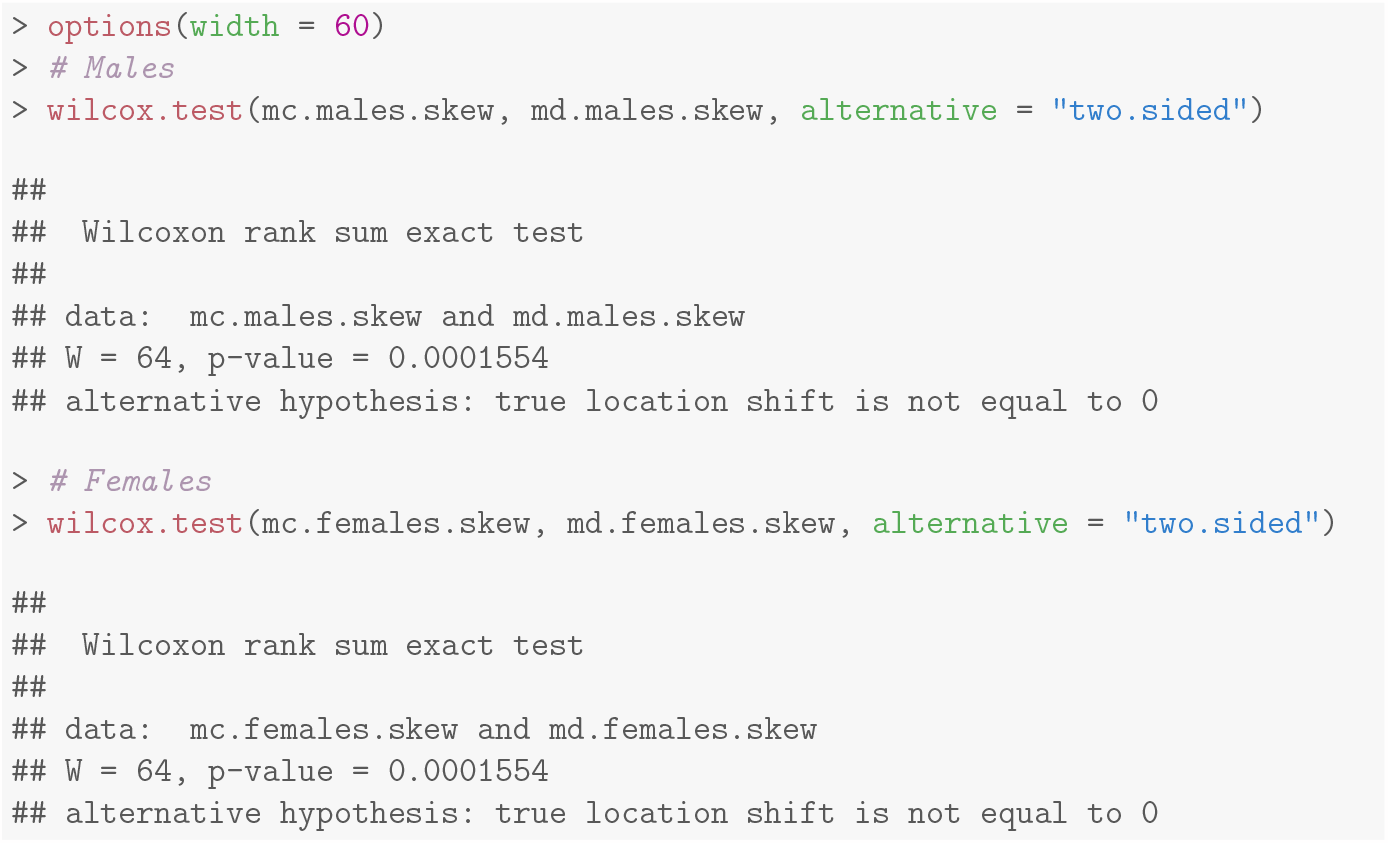

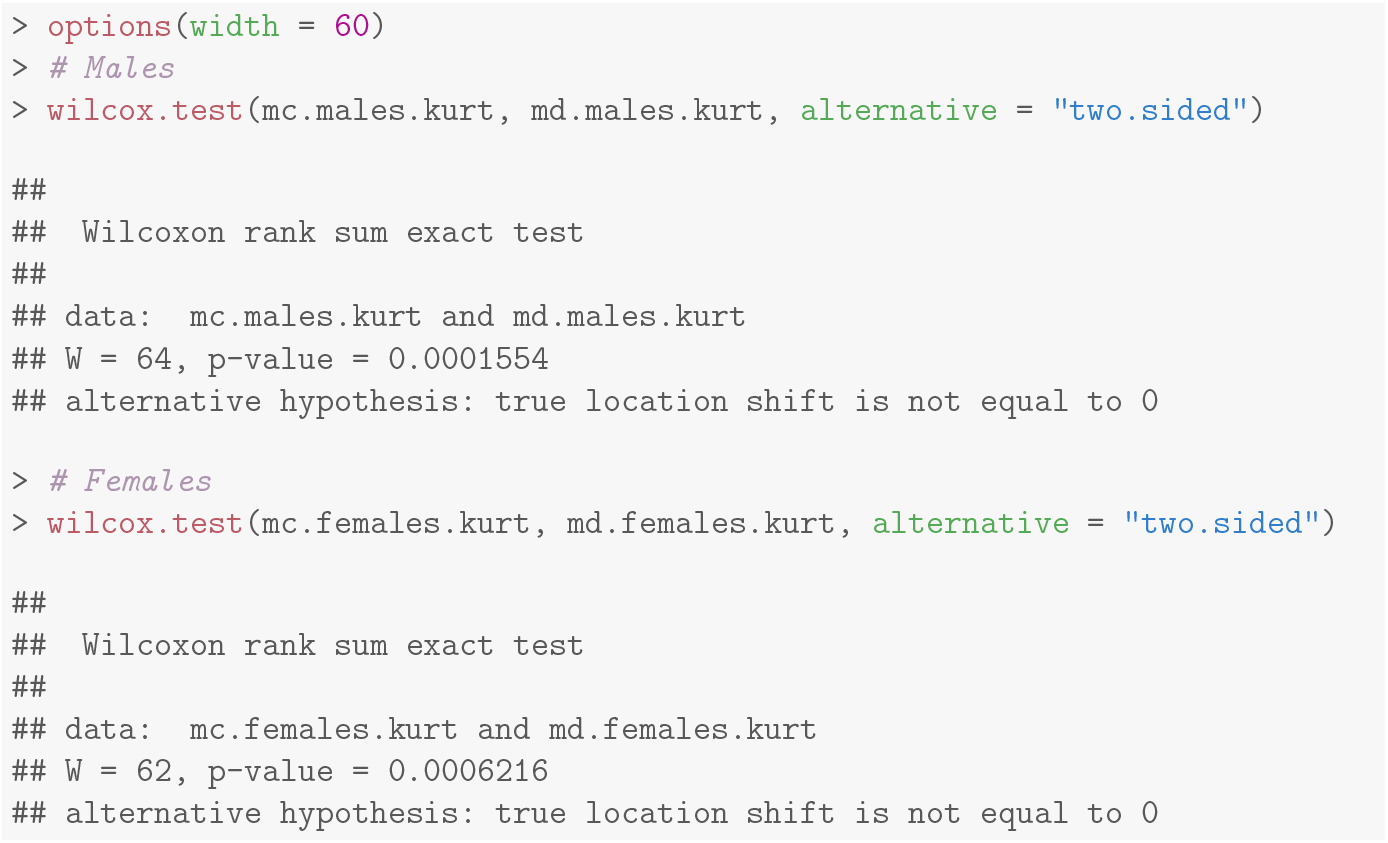

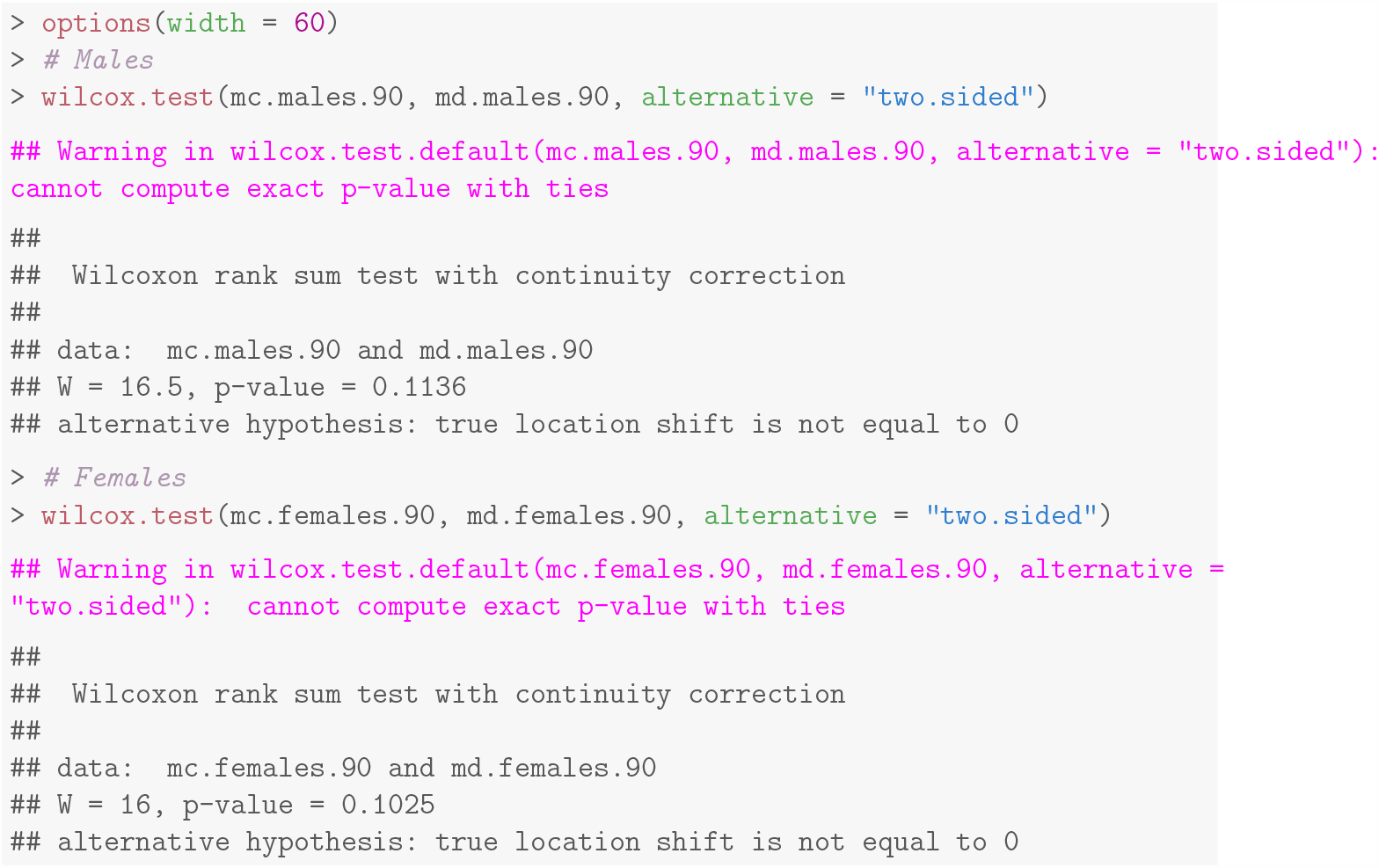

### S1.4 Goodness of fit indicators for Gamma function fits

| Popn | Sex | Block | D | KS.pval | ChiSq | ChiSq.pval |
| --- | --- | --- | --- | --- | --- | --- |
| MC | F | 1 | 0.2667 | 0.2365 | 0.1678 | 1 |
| MC | F | 2 | 0.2333 | 0.3877 | 0.0942 | 1 |
| MC | F | 3 | 0.1667 | 0.8080 | 0.1558 | 1 |
| MC | F | 4 | 0.1667 | 0.8080 | 0.1502 | 1 |
| MC | M | 1 | 0.1667 | 0.7990 | 0.0533 | 1 |
| MC | M | 2 | 0.2000 | 0.5941 | 0.0892 | 1 |
| MC | M | 3 | 0.1667 | 0.8080 | 0.0951 | 1 |
| MC | M | 4 | 0.2667 | 0.2391 | 0.1178 | 1 |
| MD | F | 1 | 0.3333 | 0.0709 | 0.2877 | 1 |
| MD | F | 2 | 0.3333 | 0.0709 | 0.4481 | 1 |
| MD | F | 3 | 0.4000 | 0.0156 | 0.4007 | 1 |
| MD | F | 4 | 0.3333 | 0.0709 | 0.2066 | 1 |
| MD | M | 1 | 0.4333 | 0.0065 | 0.1951 | 1 |
| MD | M | 2 | 0.4667 | 0.0025 | 0.1926 | 1 |
| MD | M | 3 | 0.3333 | 0.0709 | 0.2229 | 1 |
| MD | M | 4 | 0.3333 | 0.0709 | 0.2190 | 1 |

### S1.5 Kolmogorov-Smirnov tests comparing male and female kernels

| Popn | Block | D | p |
| --- | --- | --- | --- |
| MC | 1 | 0.1333 | 0.9525 |
| MC | 2 | 0.1333 | 0.9525 |
| MC | 3 | 0.1333 | 0.9578 |
| MC | 4 | 0.1667 | 0.8080 |
| MD | 1 | 0.1667 | 0.8080 |
| MD | 2 | 0.1667 | 0.8080 |
| MD | 3 | 0.1333 | 0.9578 |
| MD | 4 | 0.1333 | 0.9578 |

### S1.6 All fit parameters

| Popn | Block | Sex | k | CorrectedTheta | AIC |
| --- | --- | --- | --- | --- | --- |
| MD | 1 | M | 1.1620 | 20.3147 | -160.4278 |
| MD | 1 | F | 1.5241 | 14.0695 | -150.1206 |
| MD | 2 | M | 0.8922 | 33.9501 | -160.0143 |
| MD | 2 | F | 1.8078 | 9.9247 | -152.6892 |
| MD | 3 | M | 1.3013 | 18.5263 | -154.9708 |
| MD | 3 | F | 1.3281 | 17.9146 | -137.3899 |
| MD | 4 | M | 1.2200 | 22.3985 | -155.2544 |
| MD | 4 | F | 1.2488 | 22.2410 | -157.1292 |
| MC | 1 | M | 1.2498 | 6.4620 | -195.2162 |
| MC | 1 | F | 1.2821 | 5.9458 | -158.0278 |
| MC | 2 | M | 1.0050 | 11.8605 | -190.9815 |
| MC | 2 | F | 0.6346 | 25.2528 | -191.3530 |
| MC | 3 | M | 1.0510 | 16.5023 | -184.0834 |
| MC | 3 | F | 1.2812 | 12.0885 | -170.3489 |
| MC | 4 | M | 0.7791 | 39.5068 | -177.9508 |
| MC | 4 | F | 0.8155 | 26.9097 | -171.7862 |

### S1.7 LDD-comparing scaled and unscaled means

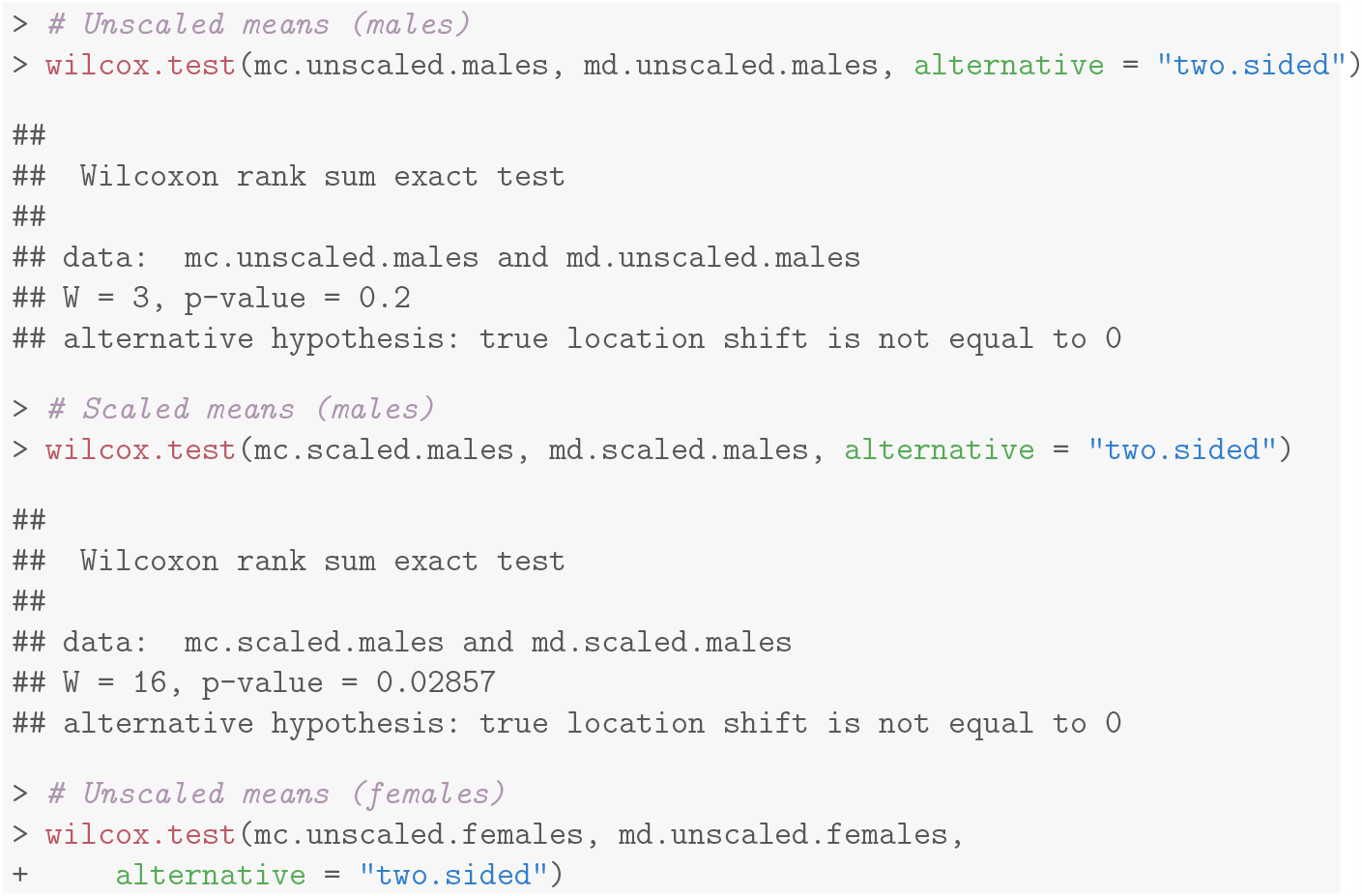

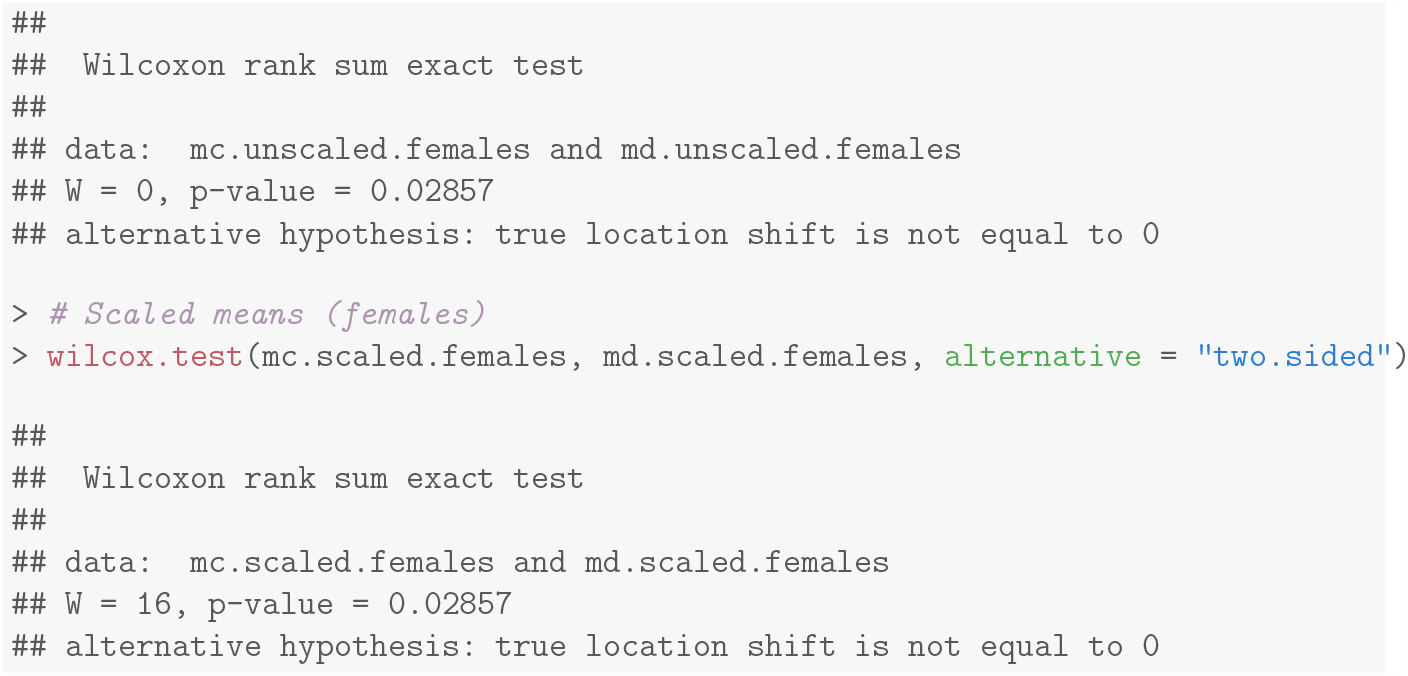

## S2 Text for

Statistical analyses for Generation 66-67

### S2.1 Propensity

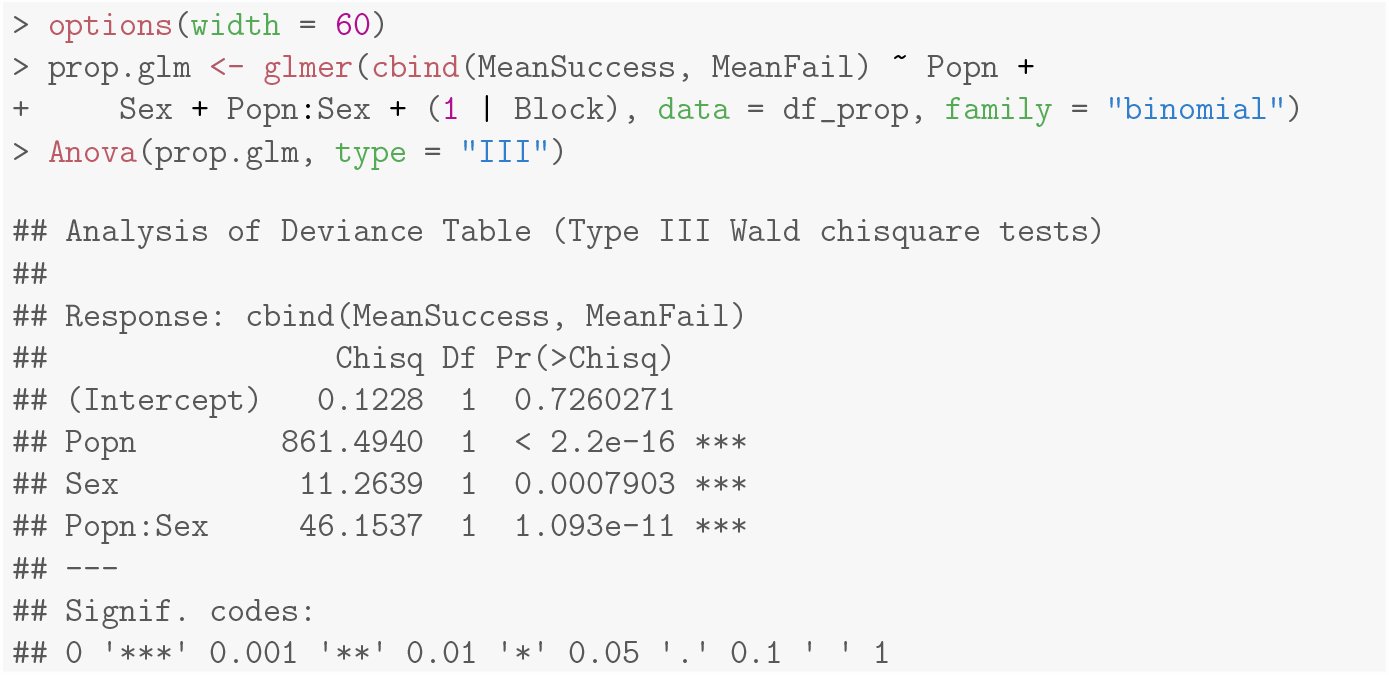

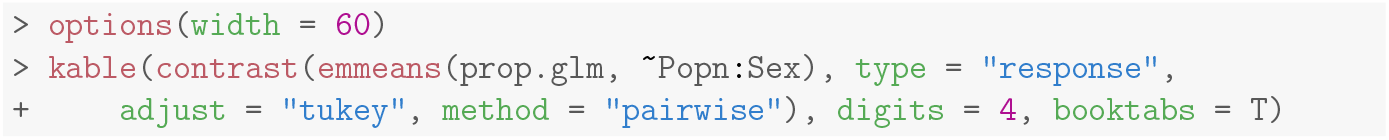

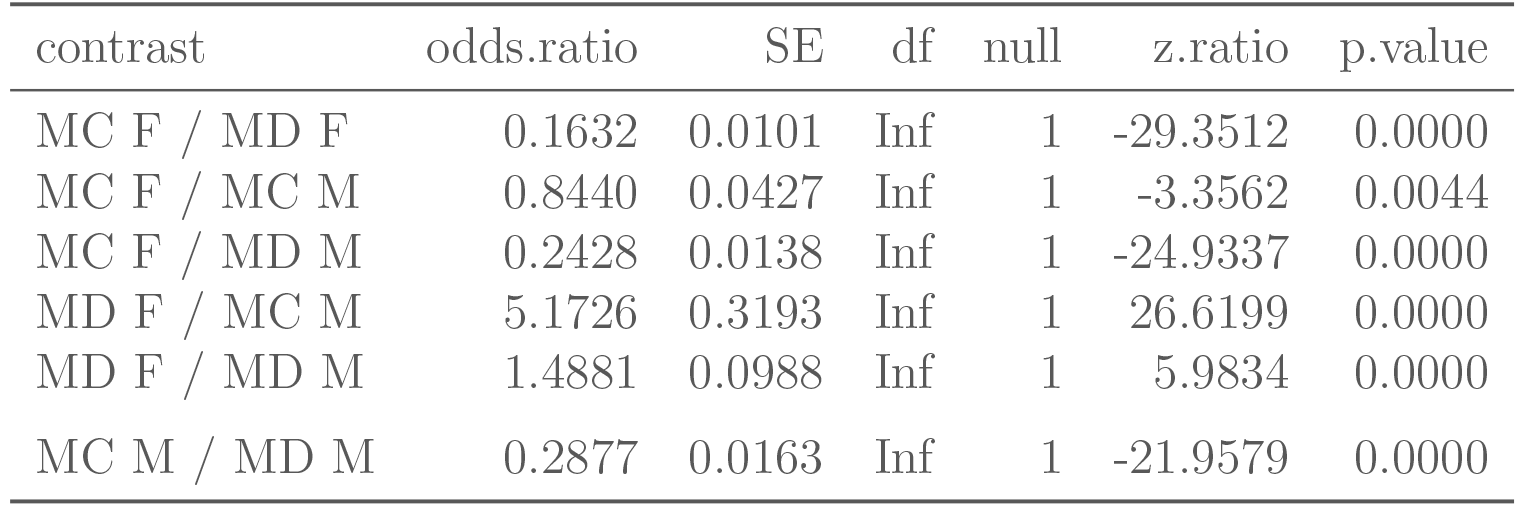

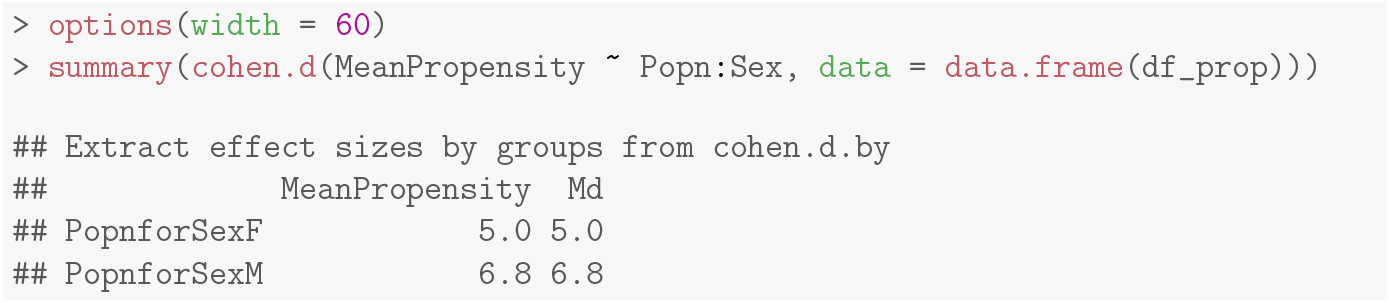

### S2.2 Ability

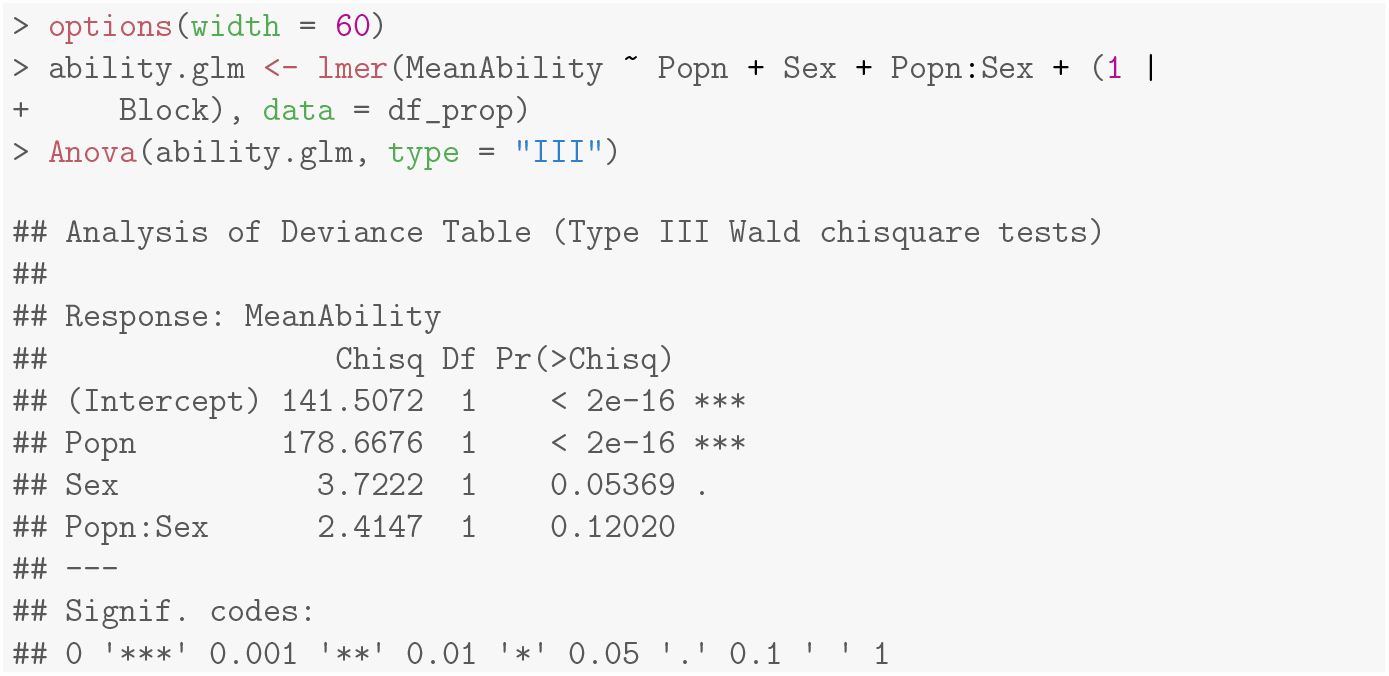

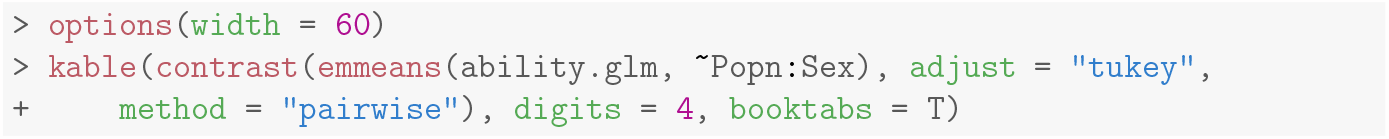

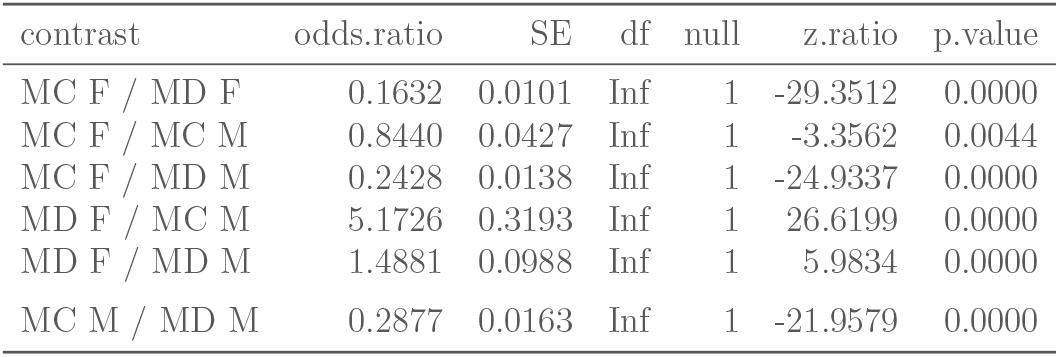

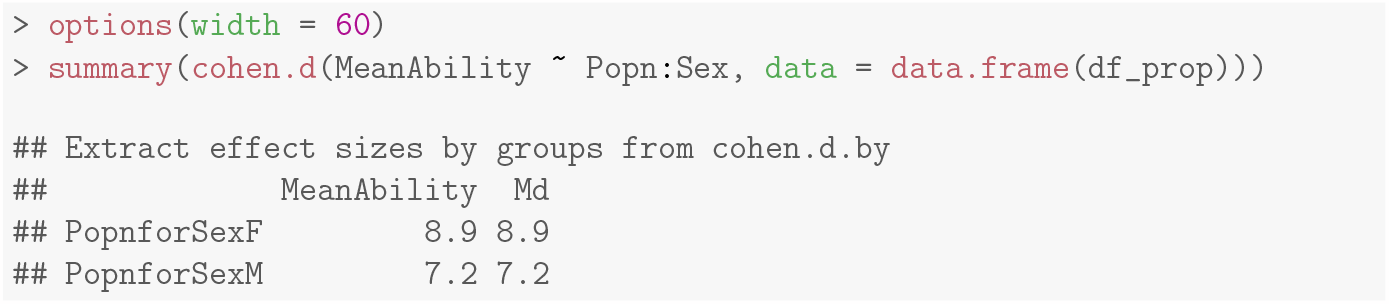

### S2.3 Higher moments of the kernel and 90th percentile

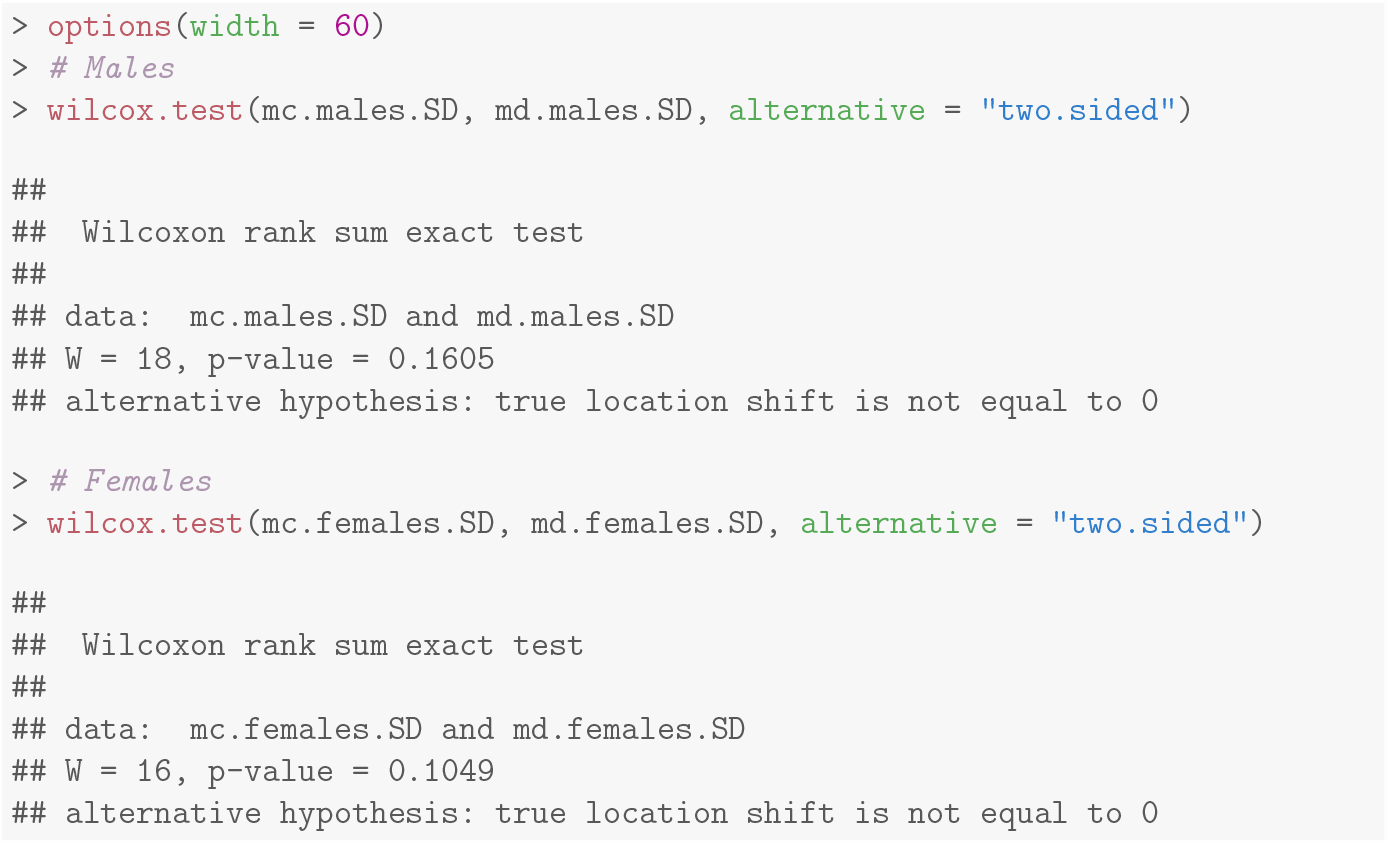

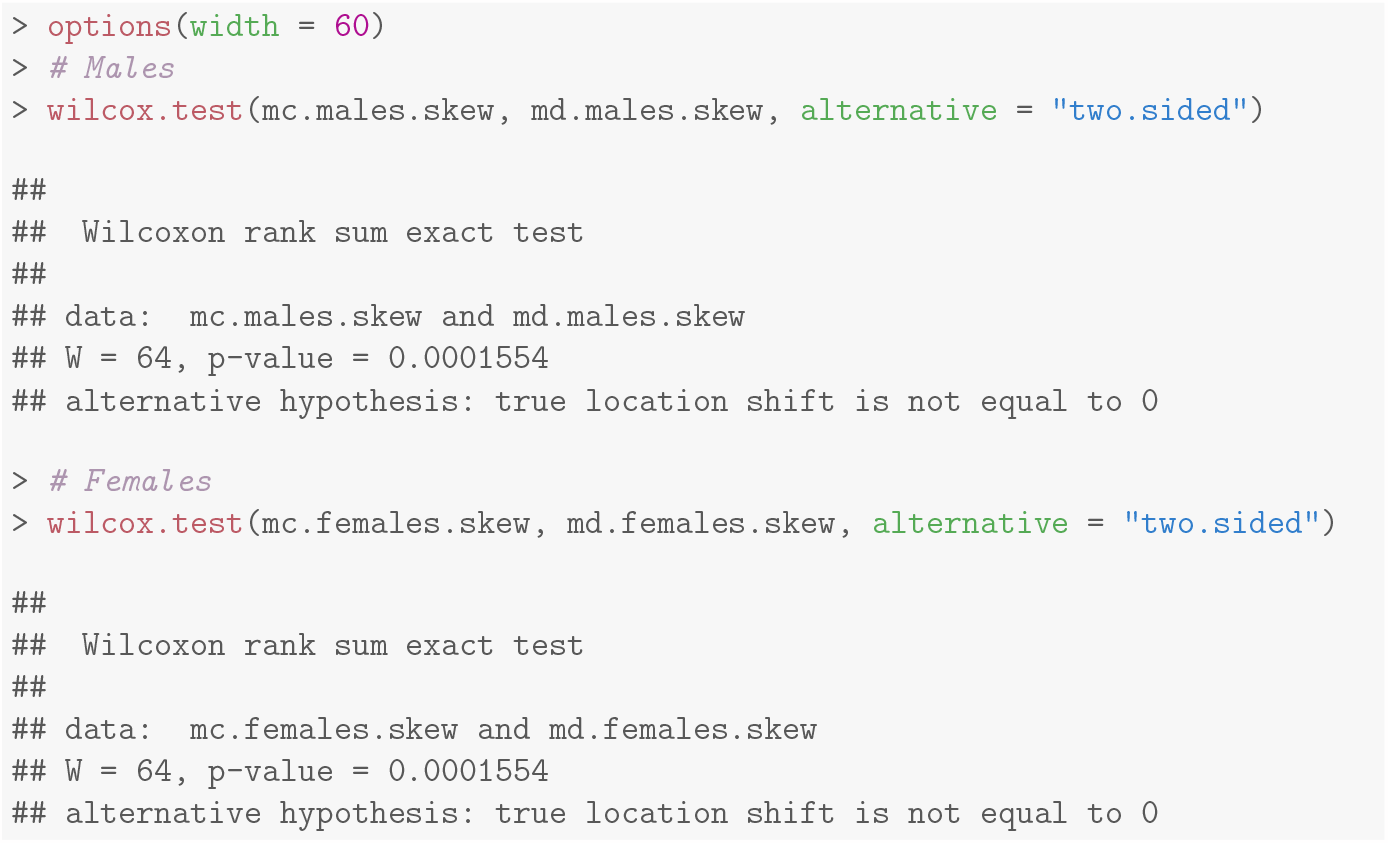

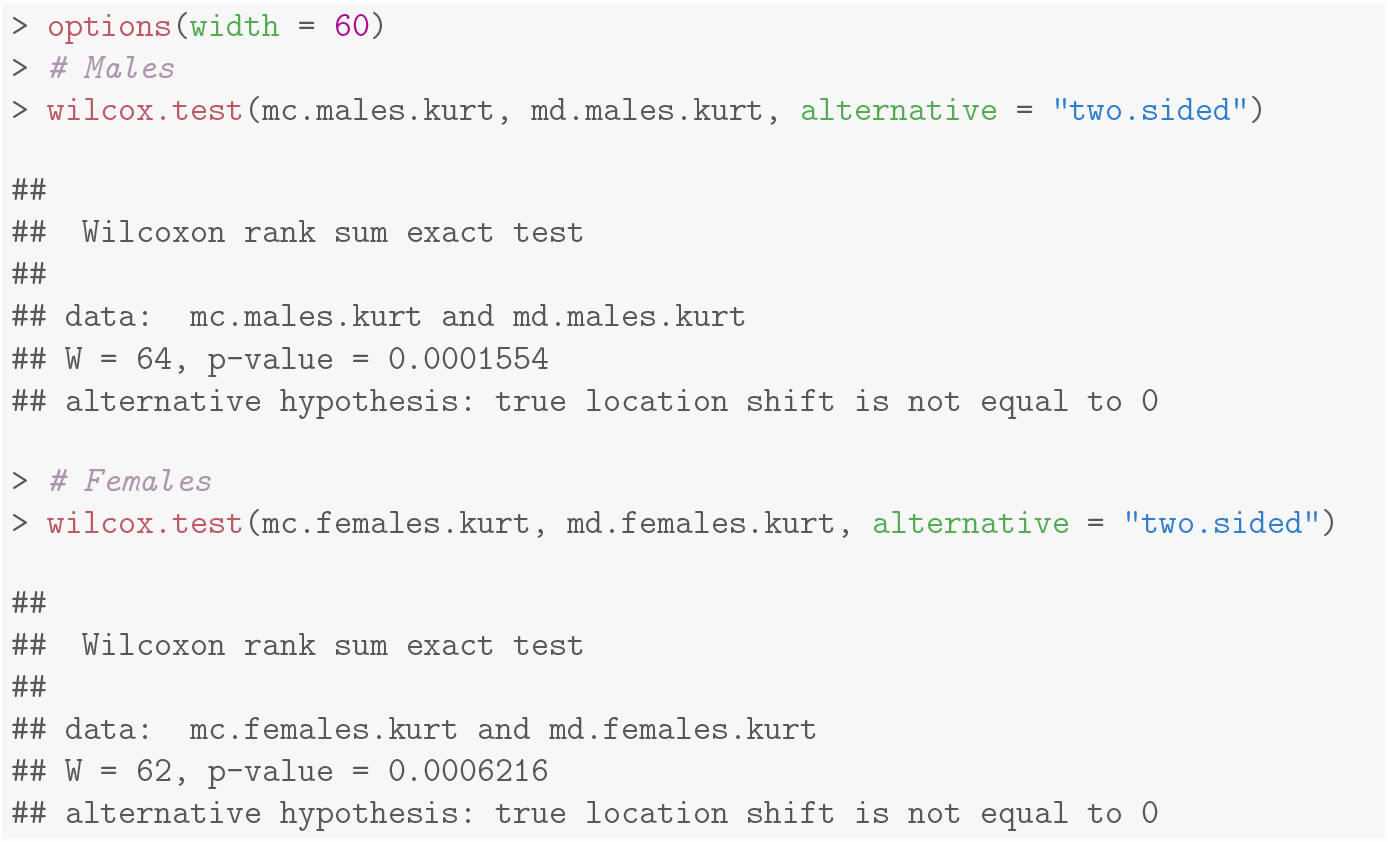

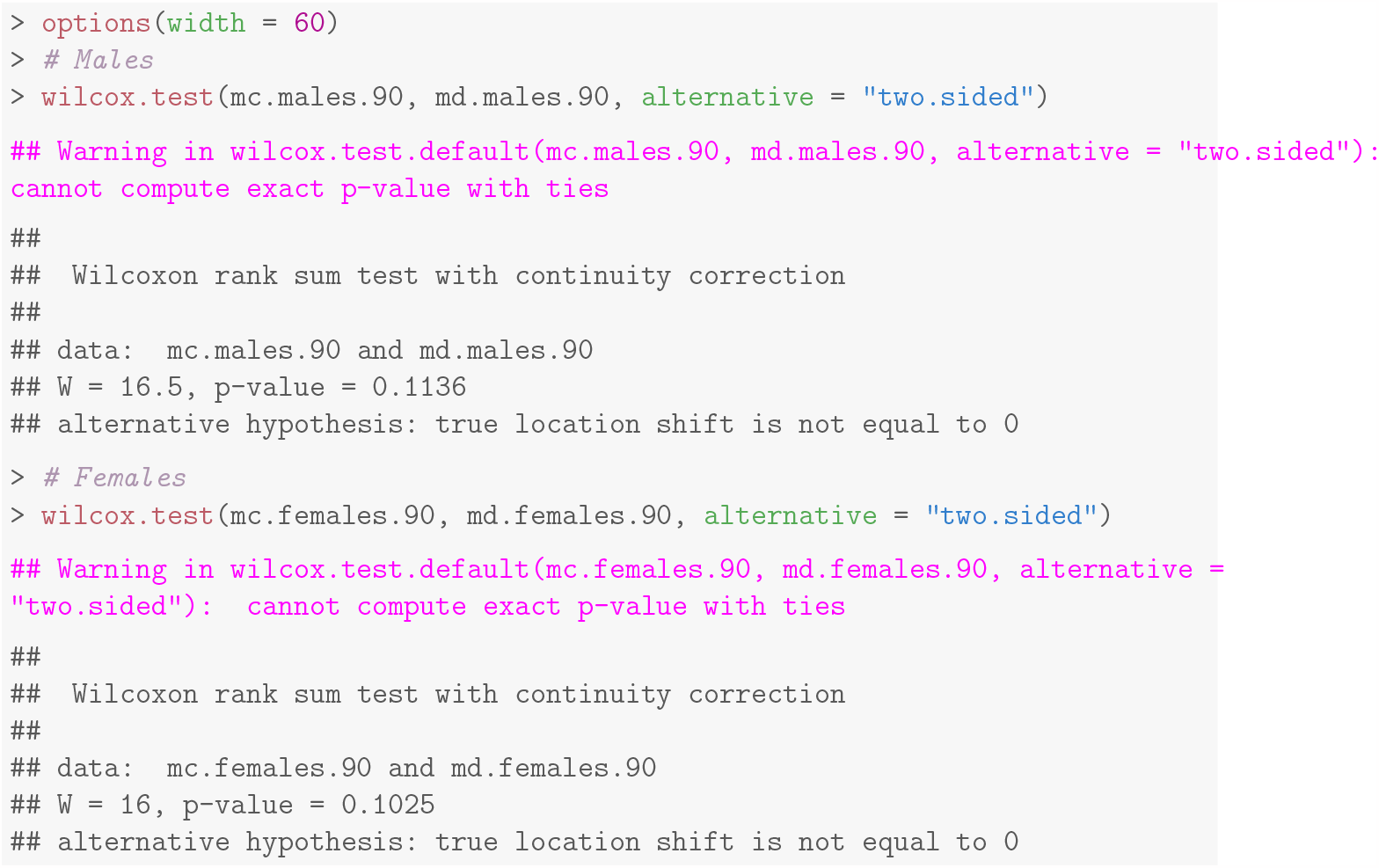

### S2.4 Goodness of fit indicators for Gamma function fits

| Popn | Sex | Block | D | KS.pval | ChiSq | ChiSq.pval |
| --- | --- | --- | --- | --- | --- | --- |
| MC | F | 1 | 0.4000 | 0.0165 | 0.0609 | 1 |
| MC | F | 2 | 0.3667 | 0.0354 | 0.1182 | 1 |
| MC | F | 3 | 0.3333 | 0.0713 | 0.0712 | 1 |
| MC | F | 4 | 0.3000 | 0.1344 | 0.0910 | 1 |
| MC | M | 1 | 0.3333 | 0.0713 | 0.1416 | 1 |
| MC | M | 2 | 0.3333 | 0.0713 | 0.0676 | 1 |
| MC | M | 3 | 0.2667 | 0.2365 | 0.0884 | 1 |
| MC | M | 4 | 0.2667 | 0.2365 | 0.0958 | 1 |
| MD | F | 1 | 0.1667 | 0.8080 | 0.0734 | 1 |
| MD | F | 2 | 0.1000 | 0.9988 | 0.4781 | 1 |
| MD | F | 3 | 0.1333 | 0.9578 | 0.2854 | 1 |
| MD | F | 4 | 0.1667 | 0.7990 | 0.6156 | 1 |
| MD | M | 1 | 0.2000 | 0.5941 | 0.0626 | 1 |
| MD | M | 2 | 0.2333 | 0.3929 | 0.0627 | 1 |
| MD | M | 3 | 0.1333 | 0.9578 | 0.0652 | 1 |
| MD | M | 4 | 0.3000 | 0.1350 | 0.1300 | 1 |

### S2.5 Kolmogorov-Smirnov tests comparing male and female kernels

| Popn | Block | D | p |
| --- | --- | --- | --- |
| MC | 1 | 0.1333 | 0.9525 |
| MC | 2 | 0.1333 | 0.9525 |
| MC | 3 | 0.1333 | 0.9525 |
| MC | 4 | 0.2000 | 0.5860 |
| MD | 1 | 0.1333 | 0.9578 |
| MD | 2 | 0.3000 | 0.1350 |
| MD | 3 | 0.1333 | 0.9578 |
| MD | 4 | 0.1667 | 0.7990 |

### S2.6 All fit parameters

| Popn | Block | Sex | k | CorrectedTheta | AIC |
| --- | --- | --- | --- | --- | --- |
| MD | 1 | M | 1.4357 | 15.9615 | -196.7267 |
| MD | 1 | F | 1.6427 | 14.0495 | -195.2812 |
| MD | 2 | M | 1.8762 | 15.1271 | -207.2036 |
| MD | 2 | F | 3.2613 | 7.5930 | -194.5058 |
| MD | 3 | M | 1.6430 | 12.4556 | -200.0810 |
| MD | 3 | F | 2.6627 | 9.3660 | -186.6777 |
| MD | 4 | M | 1.3602 | 18.5576 | -174.4135 |
| MD | 4 | F | 2.5281 | 12.0092 | -171.6101 |
| MC | 1 | M | 1.6075 | 6.2371 | -164.7455 |
| MC | 1 | F | 1.1496 | 4.8748 | -208.2715 |
| MC | 2 | M | 1.2409 | 6.9074 | -205.8391 |
| MC | 2 | F | 1.5171 | 6.3052 | -185.2442 |
| MC | 3 | M | 1.2446 | 7.7327 | -192.4412 |
| MC | 3 | F | 1.0997 | 7.3442 | -210.3603 |
| MC | 4 | M | 1.2877 | 9.0952 | -193.0083 |
| MC | 4 | F | 1.1031 | 8.2906 | -190.3597 |

### S2.7 LDD-comparing scaled and unscaled means

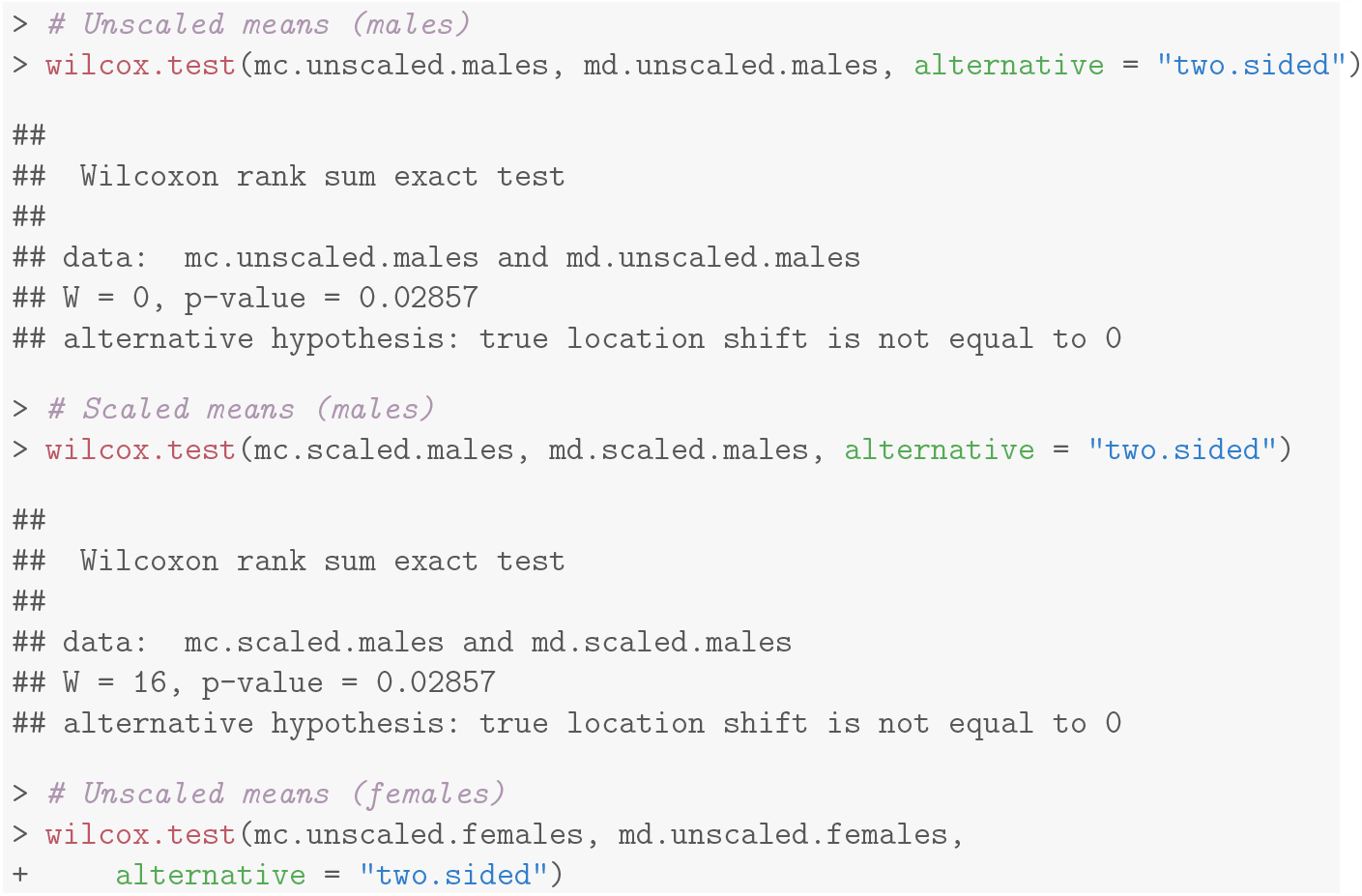

## S3 Text for

Statistical analyses for Generation 93

### S3.1 Propensity

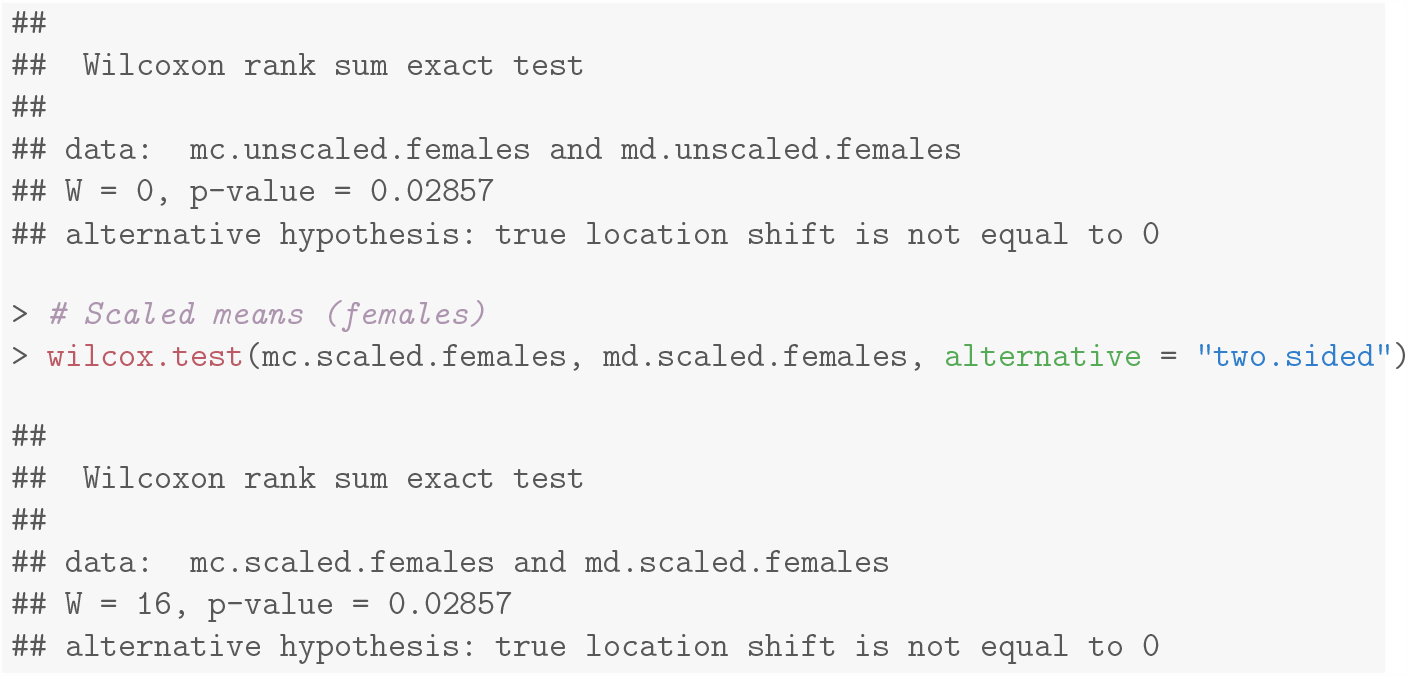

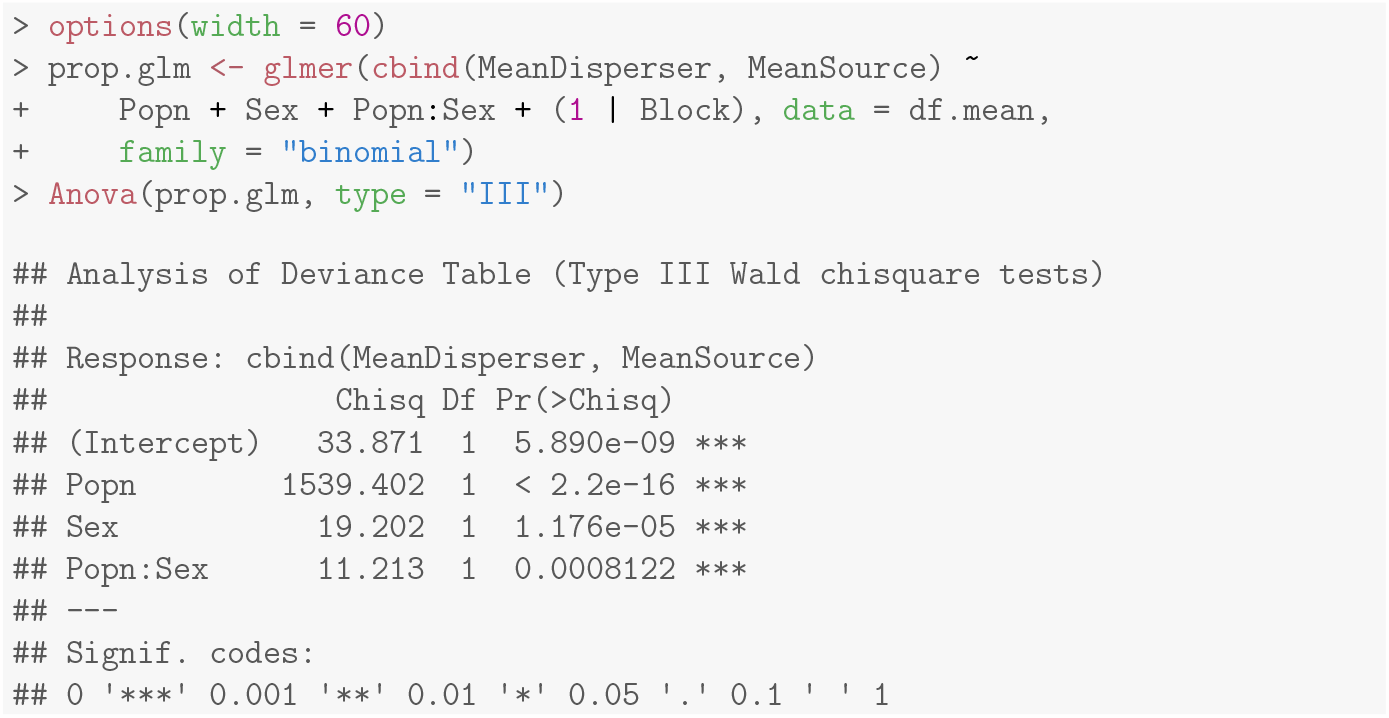

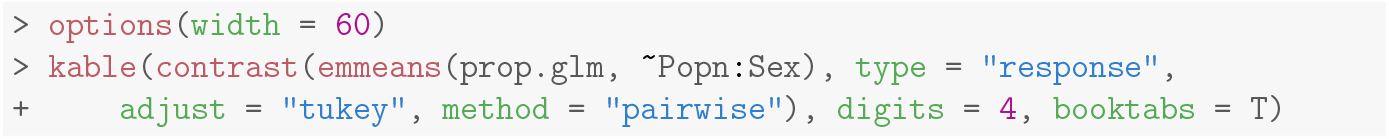

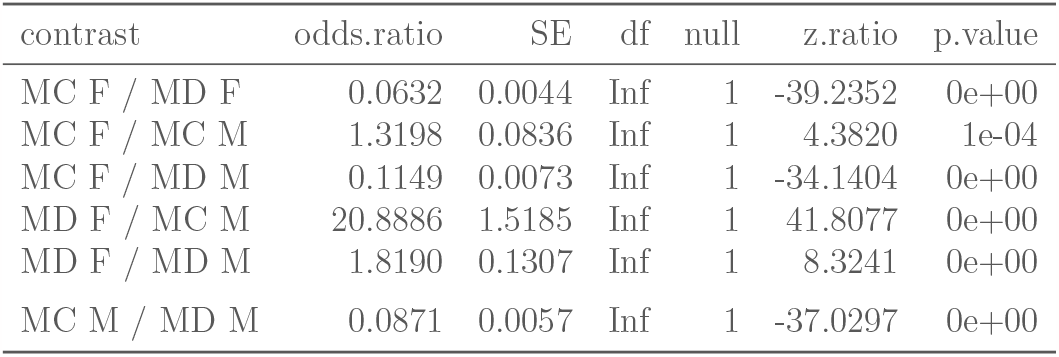

| contrast | odds.ratio | SE | df | null | z.ratio | p.value |
| --- | --- | --- | --- | --- | --- | --- |
| MC F / MD F | 0.0632 | 0.0044 | Inf | 1 | -39.2352 | 0e+00 |
| MC F / MC M | 1.3198 | 0.0836 | Inf | 1 | 4.3820 | 1e-04 |
| MC F / MD M | 0.1149 | 0.0073 | Inf | 1 | -34.1404 | 0e+00 |
| MD F / MC M | 20.8886 | 1.5185 | Inf | 1 | 41.8077 | 0e+00 |
| MD F / MD M | 1.8190 | 0.1307 | Inf | 1 | 8.3241 | 0e+00 |
| MC M / MD M | 0.0871 | 0.0057 | Inf | 1 | -37.0297 | 0e+00 |

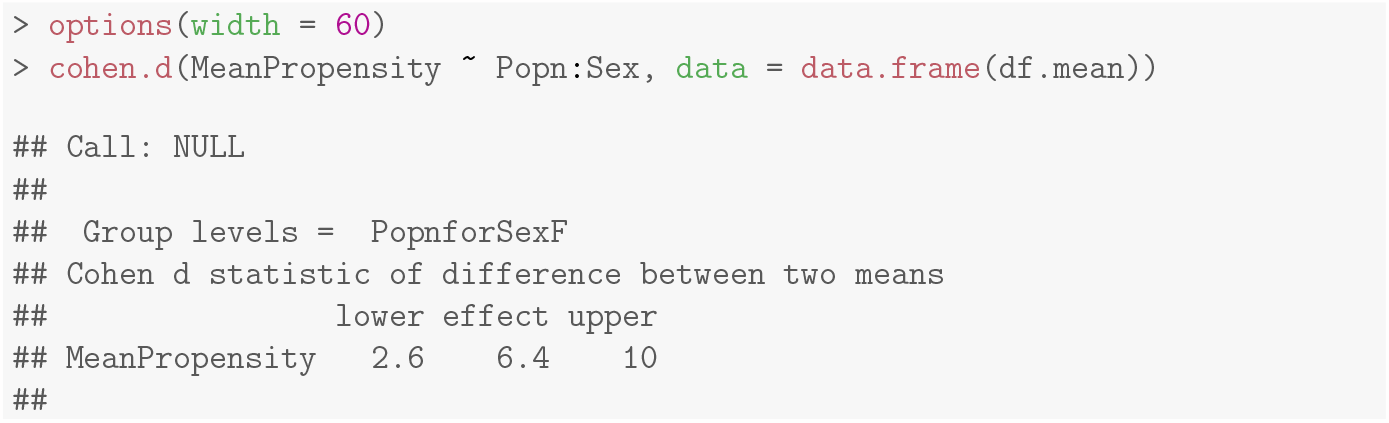

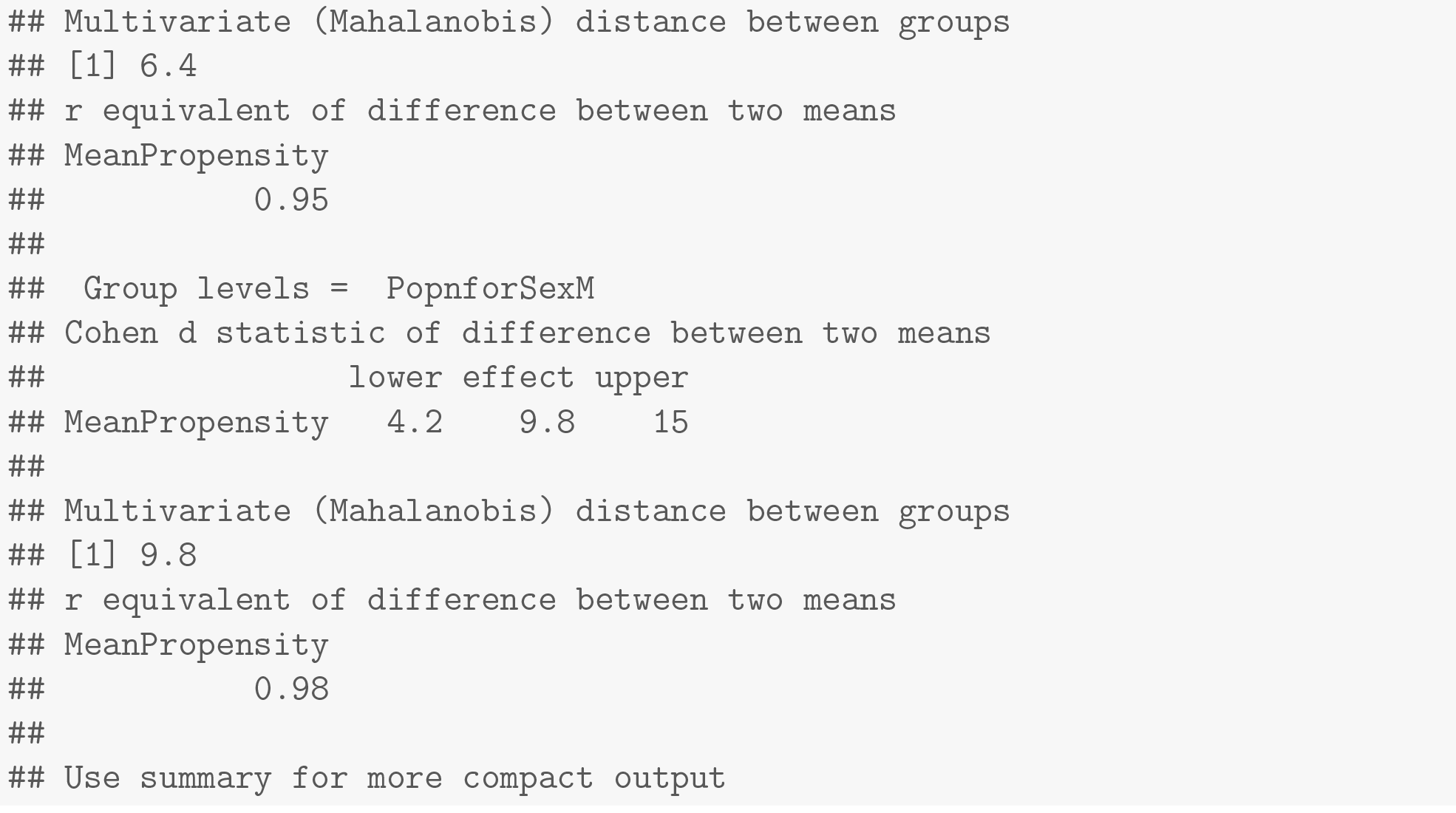

## S4 Text for

### Altered diet for larval malnutrition

We use a modified banana-jaggery medium with the amount of dry yeast powder reduced to one-third of that of the standard food. The banana-jaggery medium is a complex form of fly feed, of which most constituent parts contribute multiple kinds of complex sugars, lipids and other micronutrients (Nutrition, 1977). Unlike other simplified media (Lee and Micchelli, 2013), it is not possible to isolate a single source of carbohydrates or protein whose levels can be changed in a factorial design to produce multiple kinds of malnutrition regimes. Yeast powder is by far the simplest and most standardized component of the food as it is obtained from the same source without the kind of batch-to-batch variation that is inevitable in natural components like bananas or jaggery. Since yeast is the primary source of protein in the banana-jaggery recipe, its reduction corresponds to reduction in the amount of total available protein by about 20%, with some auxiliary reduction in carbohydrates. Standardization experiments showed a clear increase in time to eclosion as well as reduction in average dry body weight in both sexes (unpublished data) in unselected flies subjected to this malnutrition regime of one-third yeast powder, suggesting that it is strong enough to impose some amount of selective pressure over several generations.

## Supplementary Figures for

**Supplementary Figure S1:**
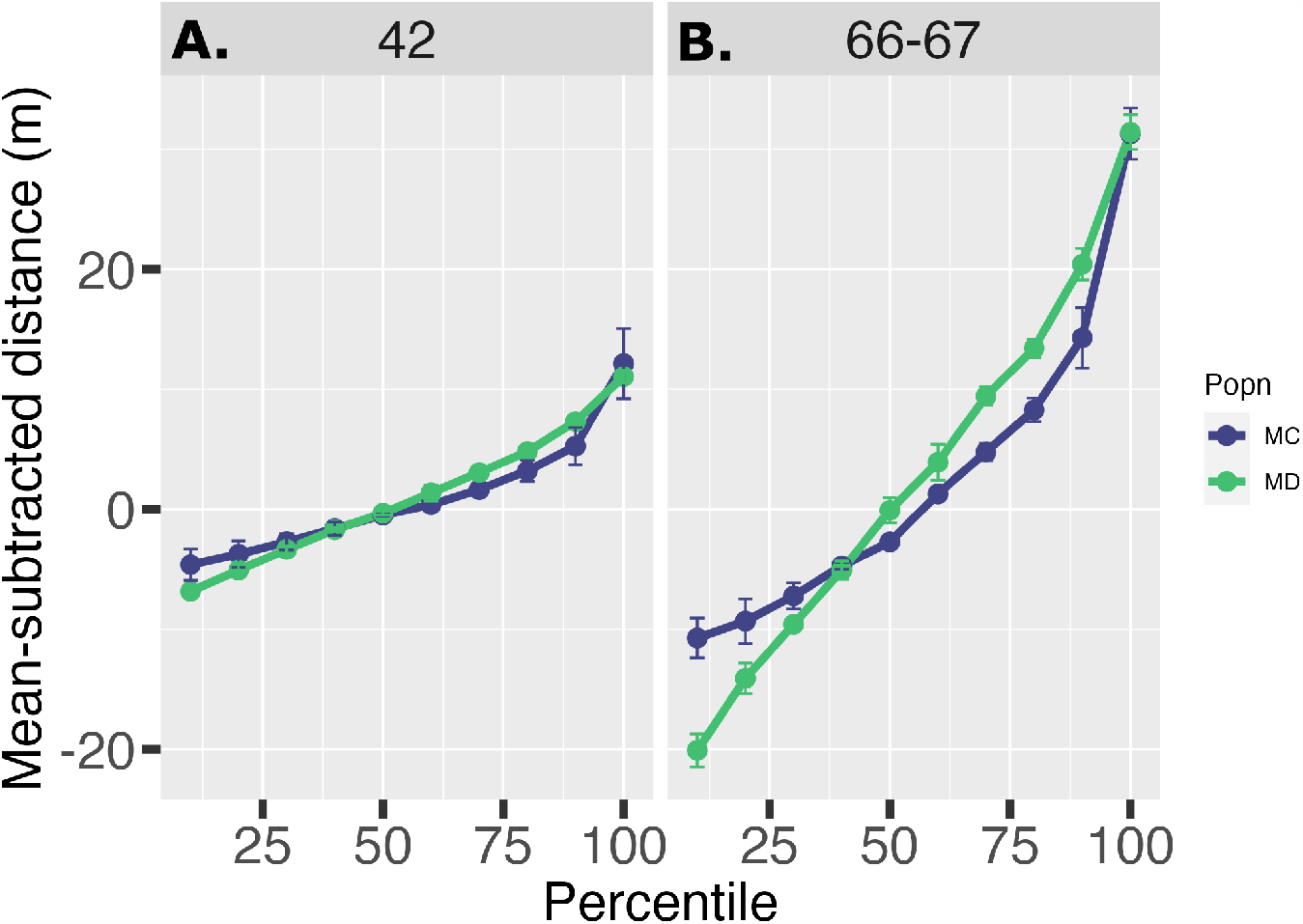
Percentiles of the mean-subtracted distances for both populations. All points are ensemble means*±*SD over the four blocks. From generation (A) 42 to (B) 66-67, spatial extent has increased for both MD and MC without a significant divergence in the frequency of LDDs, as shown by the apparent convergence of the distance percentiles beyond the 90^*th*^.

**Supplementary Figure S2:**
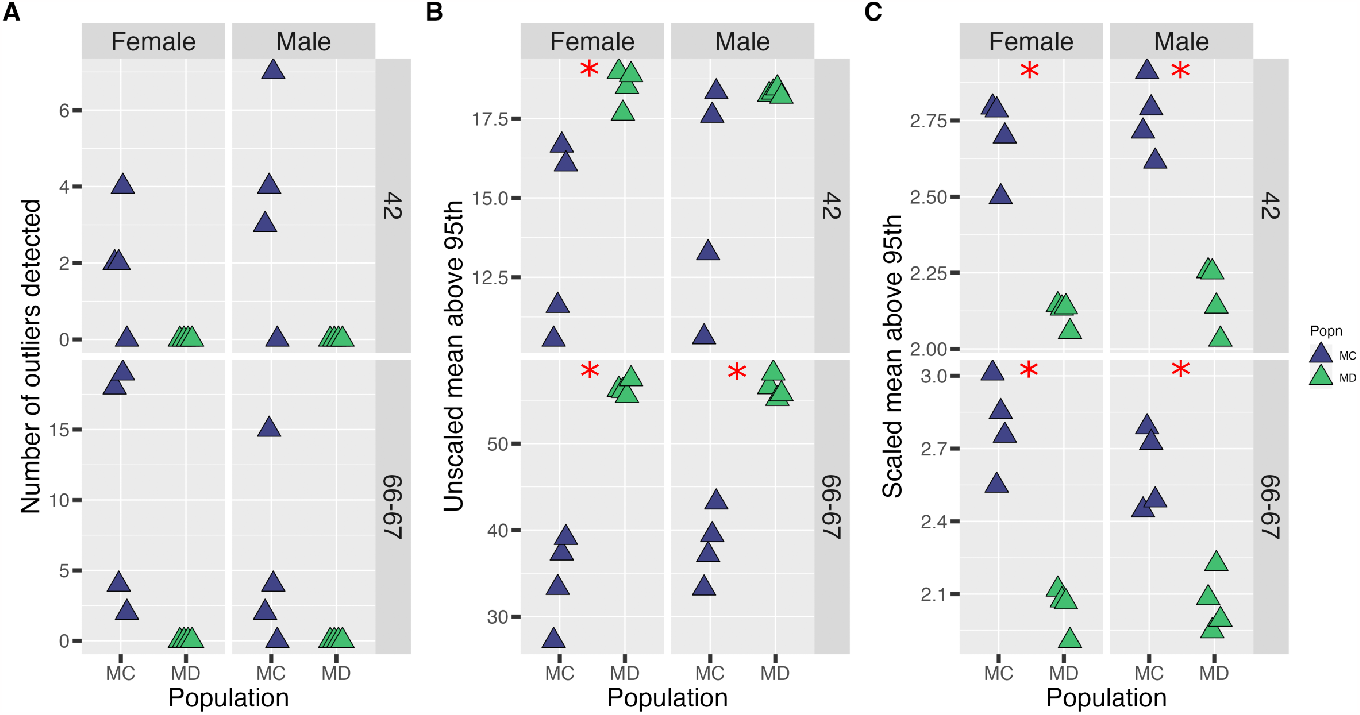
Comparison of multiple LDD measures across generations 42 (top row) and 66-67 (bottom row). (A) Number of outliers detected by the Hampel identifier with a cutoff of 2.5, (B) average value of the top 5% distances traveled (distances greater than the 95^*th*^ percentile), and (C) average of the top 5% distances, scaled by the overall average distance moved. All other specifics are the same as in Figure 5.

**Supplementary Figure S3:**
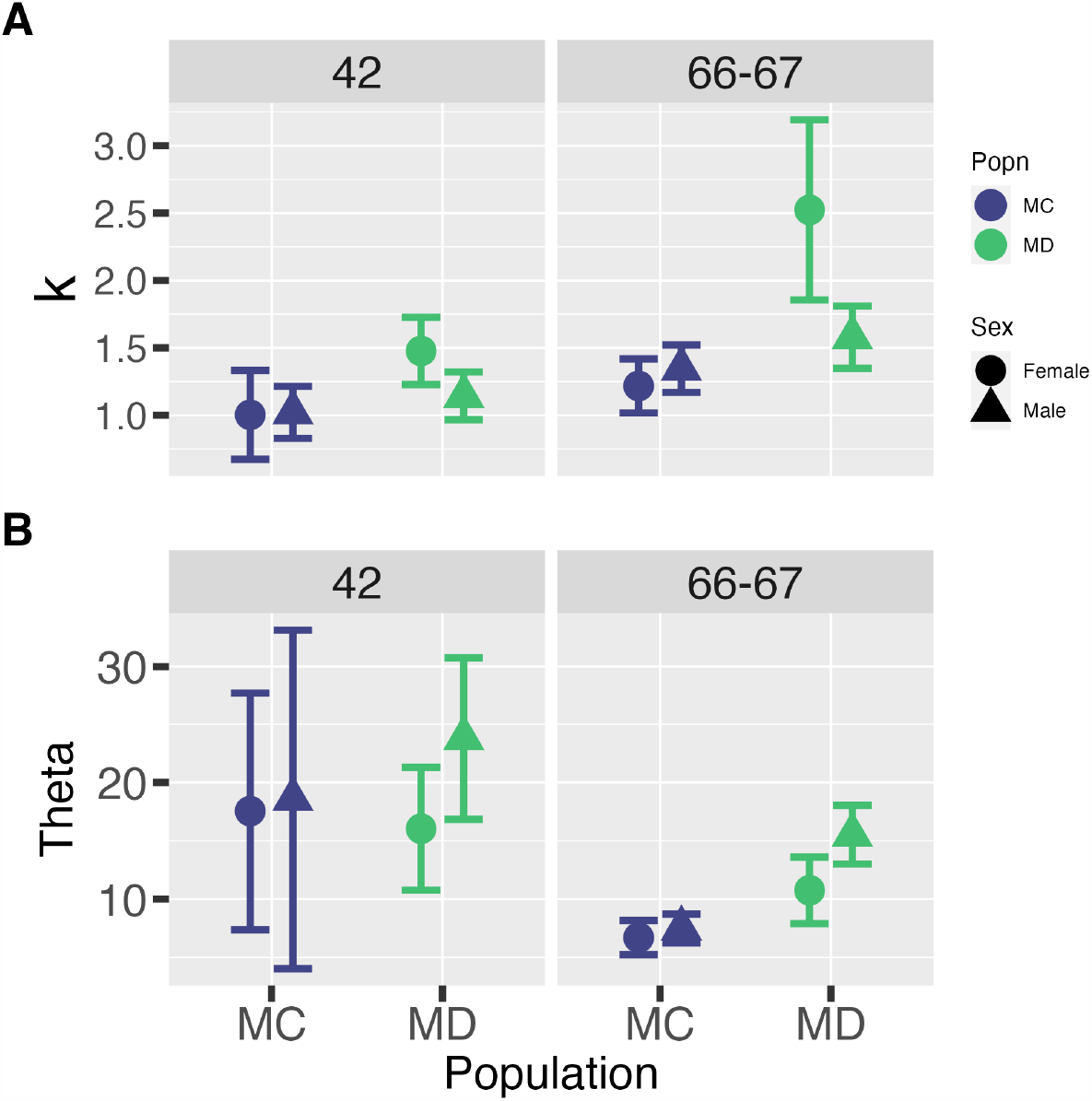
Model parameters as inferred by fitting the Gamma distribution function in Equation 1. (A) *k* and (B) *θ*^*′*^. See main text for the calculation of *θ*^*′*^. All points are ensemble means*±*SD over the four blocks. While the model parameters show some sex-specific trends, block-wise Kolmogorov-Smirnov tests comparing the frequency distributions between the two sexes within each population showed no significant differences (see S1.5 and S2.5 Text for details). It therefore remains unclear if the differences in fit parameter values between MD and MC among the sexes bear any functional significance.

**Supplementary Table S1:** Recipes of banana-jaggery medium used for stock rearing and experiments.

| **Component** | **Standard nutrition** | **Poor nutrition** |
| --- | --- | --- |
| Banana | 205g | 205g |
| Barley | 25g | 25g |
| Jaggery | 35g | 35g |
| Dry yeast powder | 36g | 12g |
| Agar | 12.4g | 20g |
| Ethanol (99.9%) | 22mL | 22mL |
| Water | 1.12L | 1.12L |
| Methyl paraben | 2.4g dissolved in 23mL of 99.9% ethanol | 2.4g dissolved in 23mL of 99.9% ethanol |

Note: The detailed procedure for food preparation is available at request.

